# RootNav 2.0: Deep Learning for Automatic Navigation of Complex Plant Root Architectures

**DOI:** 10.1101/709147

**Authors:** Robail Yasrab, Jonathan A Atkinson, Darren M Wells, Andrew P French, Tony P Pridmore, Michael P Pound

## Abstract

We present a new image analysis approach that provides fully-automatic extraction of complex root system architectures from a range of plant species in varied imaging setups. Driven by modern deep-learning approaches, *RootNav 2.0* replaces previously manual and semi-automatic feature extraction with an extremely deep multi-task Convolutional Neural Network architecture. The network has been designed to explicitly combine local pixel information with global scene information in order to accurately segment small root features across high-resolution images. In addition, the network simultaneously locates seeds, and first and second order root tips to drive a search algorithm seeking optimal paths throughout the image, extracting accurate architectures without user interaction. The proposed method is evaluated on images of wheat (*Triticum aestivum* L.) from a seedling assay. The results are compared with semi-automatic analysis via the original *RootNav* tool, demonstrating comparable accuracy, with a 10-fold increase in speed. We then demonstrate the ability of the network to adapt to different plant species via transfer learning, offering similar accuracy when transferred to an *Arabidopsis thaliana* plate assay. We transfer for a final time to images of *Brassica napus* from a hydroponic assay, and still demonstrate good accuracy despite many fewer training images. The tool outputs root architectures in the widely accepted RSML standard, for which numerous analysis packages exist (http://rootsystemml.github.io/), as well as segmentation masks compatible with other automated measurement tools.

## Background

Plant phenotyping plays a key role in plant science research, underpinning large-scale genetic discovery, and the breeding of more resilient traits [1]. This innovation makes a fundamental contribution to the push for global food security. In recent years quantitative analysis of root growth has become increasingly important as a way to explore the influence of abiotic stresses such as high temperate and drought on a plant’s ability to take up water and nutrients [2]. Segmentation and feature extraction of plant roots from images presents a significant computer vision challenge. Root images contain complicated structures, variations in size, background, occlusion, clutter and variation in lighting conditions. Fig. 1 shows an exemplar root image captured on germination paper. Even a straightforward imaging assay presents numerous challenges to a classic computer vision pipeline.

**Figure 1.**
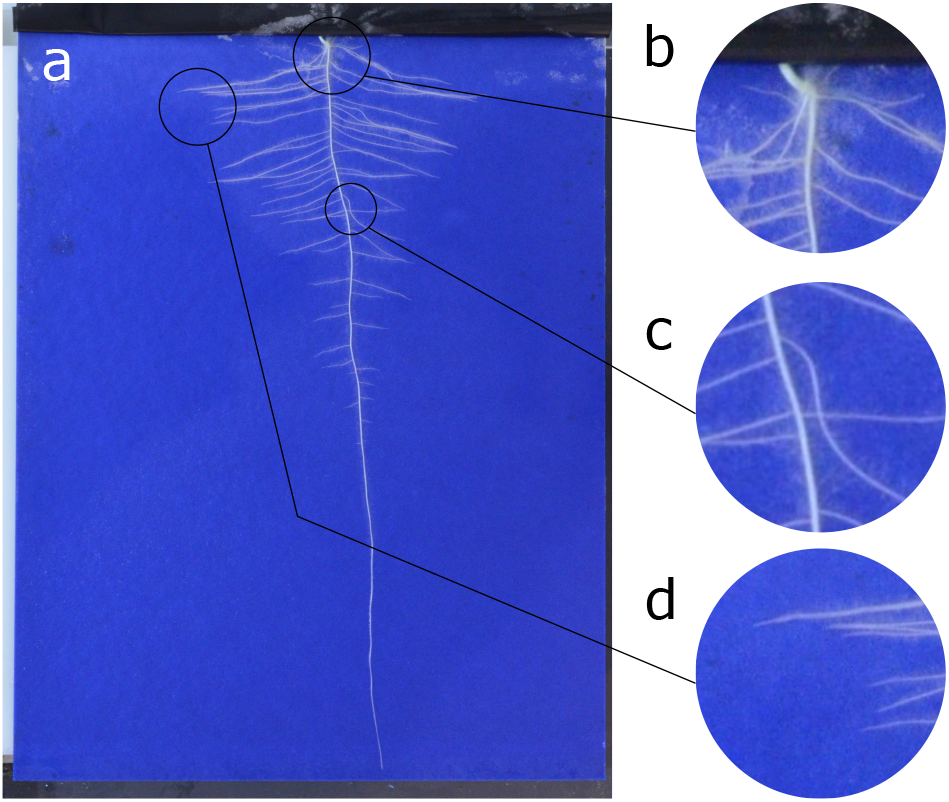
An example of the challenge root phenotyping presents for Computer Vision. a) A sample input image of a *Brassica napus* seedling grown on germination paper. This plant phenotype exhibits a single primary root and numerous lateral roots. b) Cluttered scenes make segmentation challenging. c) Complex occlusion and intersection makes extracting root topology difficult. d) Many small image features, such as root tips, occur in close proximity, making identification difficult.

In recent years machine learning has driven advances throughout many computer vision domains [3]. Indeed, much of the recent progress in plant phenotyping has also been driven by new and so-called deep learning techniques, a branch of AI, often centring around Convolutional Neural Networks (CNNs) [4, 5, 6]. The sharp increase in the availability of performant techniques in image analysis has coincided with an increase in the availability of genomic information in plant biology, providing an opportunity for robust and high-throughput solutions. The scale of the data challenge seen within plant science means that now, all but the truly fully automatic approaches will quickly become bottlenecks that hinder progress [7].

### Key Points

- A fully automatic tool for the analysis of images of root systems. Processing of one image takes between 5 and 15 seconds with no user interaction.
- Driven by a deep network in an encoder-decoder configuration, the tool outputs a complete segmentation of the root system, the location of key points, as well as an RSML description of the root architecture.
- The system can be adapted to other species and images via transfer learning, and is capable of analysing multiple plants per image.

### Analysis of Root System Architectures

In this paper, we focus on the analysis of root systems where improvements promise increases to water and nutrient use efficiency [8]. Historically, automated root phenotyping has proven challenging, due partly to the concealed nature of roots in the soil, but also to the architectural complexity and variability of root systems between species, and even individuals. Progress has been made through a combination of innovative approaches and tools [9, 10], and new imaging technologies such as X-ray and MRI (Magnetic Resonance Imaging) [11, 12].

The prevailing methodologies in root image analysis may be broadly categorised based on the level of automation they provide. Fully automated tools attempt to quantify the traits of a root system without human guidance, often through a process of image segmentation followed by post-processing. These are what might be termed ‘bottom-up’ approaches, which perform successive filtering over images in order to best distinguish between the foreground root material, and the background. Tools such as DIRT, GiaRoots, IJ-Rhizo, and EZ-Rhizo [13, 14, 15, 16], offer a familiar pipeline in which an image is first segmented into two classes, root system and background, before noise removal (such as image filters and morphology [17]) and skeletonization techniques [18] are used to clean the image. These tools then quantify the distribution of root mass within an image, providing summary statistics such as root system width, height and more complex measures such as density. Some tools, for example EZ-Rhizo, will measure root width at each location, providing more detailed analysis of the distribution of roots of different sizes.

A limitation of automated systems such as these is that errors propagate from early processing stages through to measurement. Noisy images or unexpected phenotypes will lead to errors in thresholding, which are challenging to remove and may lead to incorrect measurement of the root system. For this reason, most automated tools have placed heavy focus on cruder organ-scale measurements such as the total width of the root system, as these are most robust to small errors in image segmentation. Due to the challenge of reliably segmenting and analysing root systems automatically, many tools place strict requirements on the type of image they will analyse. RhizoScan [19], for example, offers an automatic pipeline similar to the above, based on the OpenAlea platform [20], but supports only root systems grown on Petri plates.

Beyond the problem of low-level image analysis, by framing the problem as one of identifying root pixels at a low level, these tools struggle to extract high-level root architectural information. More detailed phenotypic traits such as the number of lateral roots are out of reach of many existing tools simply because disambiguating the category of a root within a system may prove impossible in the presence of noise, especially once growth is at a mature stage where roots begin to overlap. Semantically untangling such a root system requires a higher-level understanding of the image than pixel-based processing methods provide.

Manual root analysis tools such as ImageJ’s polyline function [21] and DART [22] offer an entirely different approach. They place reliance on an expert human annotator to successfully identify the structure of the root system by asking the expert to label each root by hand. The advantage here is that if sufficiently well trained, an annotator could conceivably reconstruct an entire root system, using their advanced knowledge to clear up disambiguation in cluttered areas of the image. The obvious drawback to this approach is that this is an extremely time-consuming process. In practice, many experiments will therefore have to severely limit the number of measurements captured per image, such as by focusing on primary root length, to bring the time required into a reasonable range. Some tools, for instance RootScape [23], have been designed with this in mind, requiring that a user highlights only 20 key landmarks on a root system. These landmarks are then used to explore phenotypic differences between genotypes via principal component analysis. In those instances where detailed analysis is required, the burden on annotators is huge, and the cost of mistakes may be high. Outside of plant science, obtaining cheap and efficient annotation has become a widely researched topic in and of itself [24, 25]. In plant science, noisy and low-cost annotation may not be acceptable, depending on the experimental requirements, and ultimately offers few benefits over the automated tools described above.

Alongside the development of manual and automated tools, a selection of widely-used semi-automatic tools have been released. These approaches aim to bridge the gap between speed and accuracy, offering a compromise acceptable for many use-cases. Tools such as RootReader [26] perform a similar automatic function to the tools above, but provide the user with the ability to manipulate some of the output to correct mistakes. Most of the tools in this category are not bottom up, and instead model the root system in some way, guided by the user, in order to better understand the image on which they are run. Smartroot [10], a plugin for the popular ImageJ tool [21], operates by tracing along each root in a guided way, at each step searching for the optimal direction in which to travel based on the current orientation of the root at that point. Smartroot is semi-automatic, with initiation of roots and correction of errors often requiring human intervention. Nevertheless, with some user effort Smartroot can potentially be used to reconstruct full root system architectures. RootNav [9], a precursor to the work presented here, offers a point-to-point path search between labelled seed locations and root tips. Images are first segmented into background/foreground classes, before a user is required to label root tip and seed locations. Shortest path search is used to trace between key organ landmarks, resulting in a complete reconstruction of the root system. However, RootNav does not include a reliable method for *detecting* seeds and root tips (the user must perform this step), nor is the segmentation step robust to image noise. This means that significant user interaction is still required to guide the software, but as with Smartroot, the output is a full and architecturally correct root system architecture. Many tools that are able to output root system architectures have been adapted to provide output in the popular RSML (Root System Markup Language) format [10]. RSML is an XML-based standard for the sharing of root system architectures, including information on geometry, and relative position within the system. Numerous tools exist to read and write RSML files, allowing customised pipelines between tools, and the ability to decouple the image analysis from the ultimate measurement of traits, as well as view the final architecture labelling.

### Deep Learning for Root Systems

The prevailing methodology when working with images in deep learning is the Convolutional Neural Network (CNN). CNNs improve upon traditional machine learning via their ability to learn not only solutions to problems, but also the most effective way in which to transform data to make this goal easier. This representation learning provides CNNs with unparalleled discriminative power, and has seen them quickly move into a dominant position within the field of computer vision [3]. A CNN is a layered structure that performs successive image filtering operations that transform an image from a traditional RGB input, into a new feature representation. This transformation is learned during training, and provides the final layers of the CNN with the best possible view of that data from which to base decisions. The deeper into a CNN data flows, the more abstracted and powerful the representation becomes. While the initial layers may compute simple primitives such as edges and corners, deeper into the network feature maps may highlight groups of primitives. Deeper still, feature maps may contain complex arrangements of features representing realworld objects [5]. These features are learnt by the CNN training algorithms, and are not hand-coded, meaning that with sufficient training data any number of different problems can be addressed. Within the Biosciences, such networks have been used to perform a variety of tasks ranging from classification, assigning discrete labels to images and objects [27] through to regression problems; that is, of directly predicting values [28]. For root systems, [5] used a deep classification network to scan an image for probable root tip locations in 32×32 pixel tiles. Despite promising results, the drawback of this approach is that using a small field of view, customarily called a *receptive field* within the machine learning literature, is computationally less efficient, and may cause produce additional false positives where the small field of view is not sufficient to distinguish true roots from image noise. This system also only currently detects root tips, which means more complex traits involving other organs cannot be computed.

### Image Segmentation and Feature Localisation

The measurement of complex phenotypic traits requires analysis at a finer scale than that of whole-root-system traits, but sensitive to more than only a small selection of plant features such as just root tips. To address this, the research community has begun to move towards networks that output a richer array of information. Recent work has been based around newer CNN designs in what we term an encoder-decoder configuration, aimed at segmentation of images, or the location of key feature points. Traditional CNNs perform spatial downsampling such that by the end of the network, features spatially correspond to the entire image, i.e. they have lost location resolution. This is ideal for classification tasks, where a decision must be made on an image scale. This is not appropriate, however, for situations where a 2D segmentation result is required. Encoder-decoders therefore upsample again from the feature space, back into a spatially high-resolution image (Figure 2). This process can be thought of as combining a CNN with a second, reversed CNN that learns to produce images once again, these images might be trained to predict the locations of objects, or to segment pixels into background and foreground classes. Encoder-decoders are being used in plant science to, among other tasks, segment plant shoots [29, 30], other plant organs [31], and fill gaps in rhizotron images of root systems [32]. The work in [5] first introduced the concept of heatmap regression to the plant phenotyping domain, in which a segmentation output is replaced by a heatmap showing likely target locations. Our development in this paper combines both of these approaches, simultaneously segmenting a root system, *and* predicting the likely locations of root tips and seeds.

**Figure 2.**
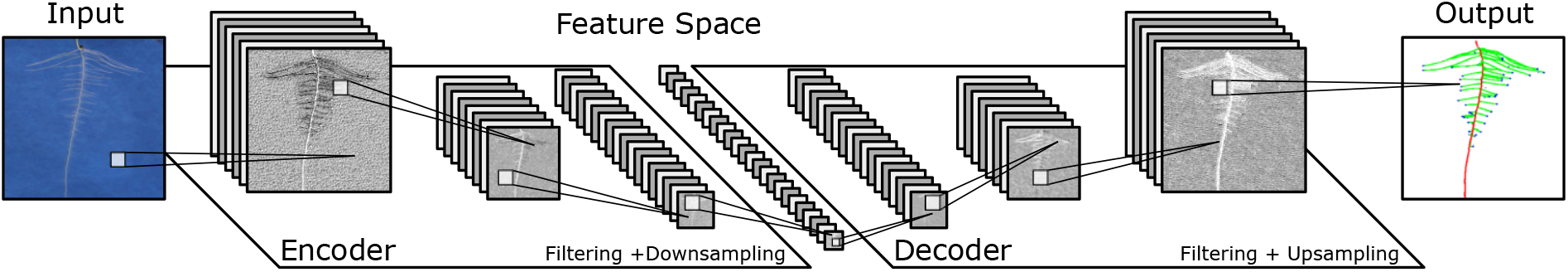
A simplified example illustrating the major components of a CNN in an encoder-decoder configuration. The encoder performs a combination of filtering operations including convolutional filters, spatial downsampling, and normalisation. These layers convert the original image into a high-dimensional feature space, but with very low spatial resolution. The decoding network performs similar layer operations, but replaces downsampling with upsampling to return the feature representation back into a spatially high-resolution image.

### Automated Root Phenotyping

We present here a new tool for the automatic analysis of root systems which is designed to work across a wide variety of plants and imaging conditions. Our pipeline is driven by a deep encoder-decoder network, similar to that presented in [5] but adapted to handle higher-resolution images. The network is trained to simultaneously segment root material, classify root type, and locate key features from which root geometry can be derived. To our knowledge this is the first use of deep learning to perform multi-task segmentation and localisation in plant phenotyping. The output of the network is refined using an A* shortest path algorithm to determine the most likely path of each root, connecting located second order roots to appropriate first order roots, and first order roots back to the seed location. Full root geometry is extracted per plant, and is robust to multiple plants and highly varied architectures. The tool outputs the standard RSML format [33], widely supported by the community, from which RSA traits may be derived. The tool also outputs the underlying segmentation masks for first and second order roots, from which global traits may be derived. An overview of the tool can be seen in Figure 3. The system first performs pixel-wise segmentation of the image, and heatmap regression to locate key features, it next extracts the root topology via a series of guided shortest path searches, before finally extracting the entire root architecture into a portable RSML format.

**Figure 3.**
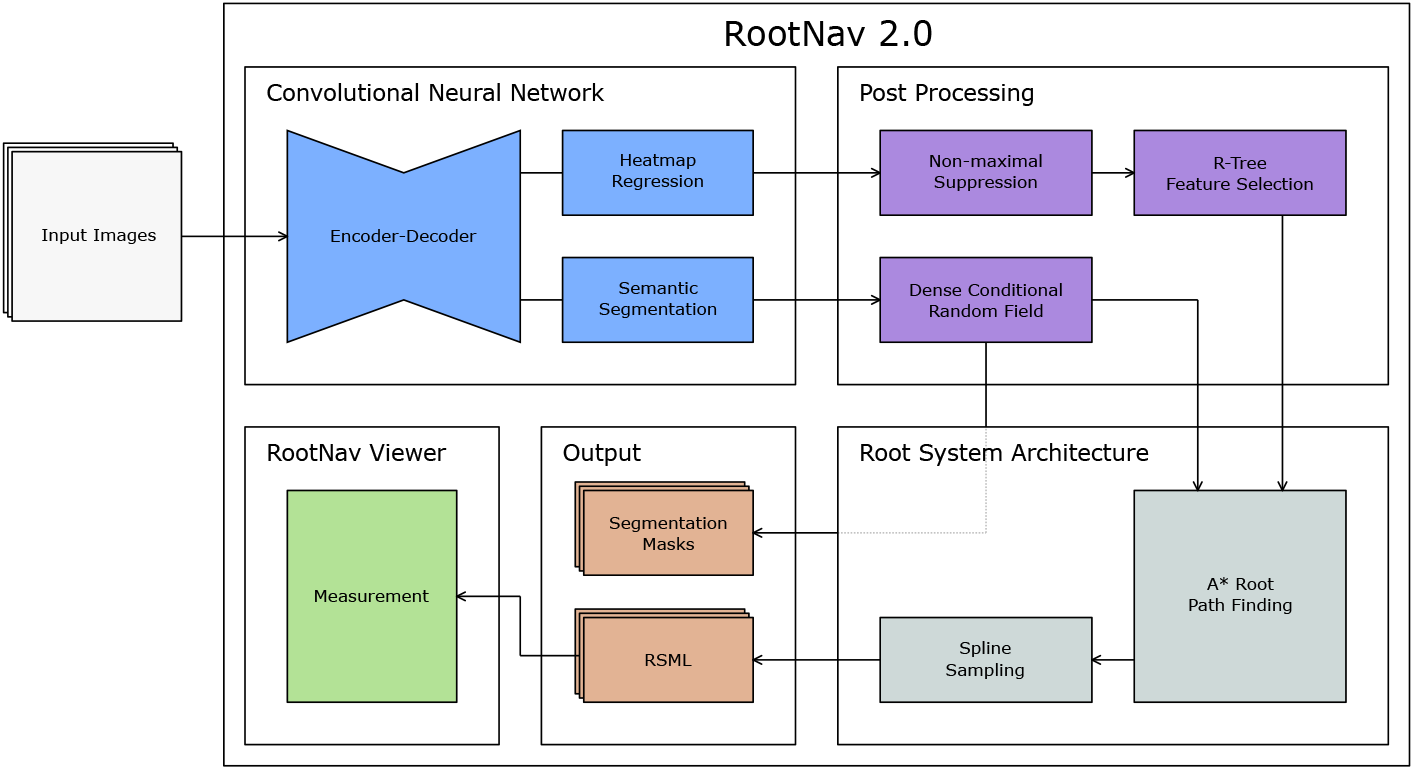
An overview of RootNav 2.0. The input enters a CNN that performs both segmentation of the root structure, and localisation of key points. These are post-processed to extract information for a path finding algorithm. A* search then extracts likely paths taken by each root, generating an entire architecture for an arbitrary number of plants in an image. All roots are resampled as smooth splines, before all topology and geometry is output into an RSML file. Segmentation masks for first and second order roots are also saved.

We first demonstrate the performance of the tool on a large wheat dataset grown on germination paper. We perform a quantitative comparison with traits measured using the original semi-automatic RootNav tool [9], hereby referred to as RootNav 1.0, in which an expert performed detailed manual intervention to ensure accuracy. We next demonstrate the ability of RootNav 2.0 to adapt to new image types with a much smaller training set. We retrained the network on 200 images of Arabidopsis grown on agar plates, in which up to five plants appear per image. We again compare quantitatively against human labelled images generated using RootNav 1.0. Finally, we transfer learn once more using an even smaller, rapeseed dataset, comprising only 91 training images. Beyond accuracy measures, we have assessed our system’s performance in terms of inference time and resource efficiency, to provide a comparative analysis of user burden for root architecture analysis. The trained networks, tool and all training datasets will all be made publicly available.

## Data Description

### Primary Dataset

Our primary dataset is composed of images of Wheat (*Triticum aestivum* L.) seedlings totalling 3,630 images of 1900×2000 pixel resolution. Images include those released in [5], plus additional images captured using the same methodology. Images were captured as per [34]; seeds were sieved to uniform size, sterilized, and pre-germinated before transfer to growth pouches in a controlled environment chamber (12-hour photoperiod: 20°C day, 15°C night, with a light intensity of 400 µ*mol m*^−2^ *s*^−1^ PAR). After 9 days (with plants at the two-leaf stage), individual pouches were transferred to a copy stand for imaging using a Nikon D5100 DSLR camera controlled using NKRemote software (Breeze Systems Ltd, Camberley, UK). Ground truth annotations for all plants were obtained using the original RootNav 1.0 software [9], and stored in RSML format [33]. Each annotation was provided by an expert user, and as we intended to use RootNav 1.0 as a quantitative baseline for accuracy emphasis was placed on accuracy over speed during this process.

Ground truth images for network training and validation were generated from these RSML files by rendering appropriate segmentation masks and heatmaps. The dataset was split into training and validation sets totalling 2,864 and 716 images respectively. An additional 50 images were held back as a final testing set. More details on this methodology may be found in the Methods section. Example images can be found in Figure 4a.

**Figure 4.**
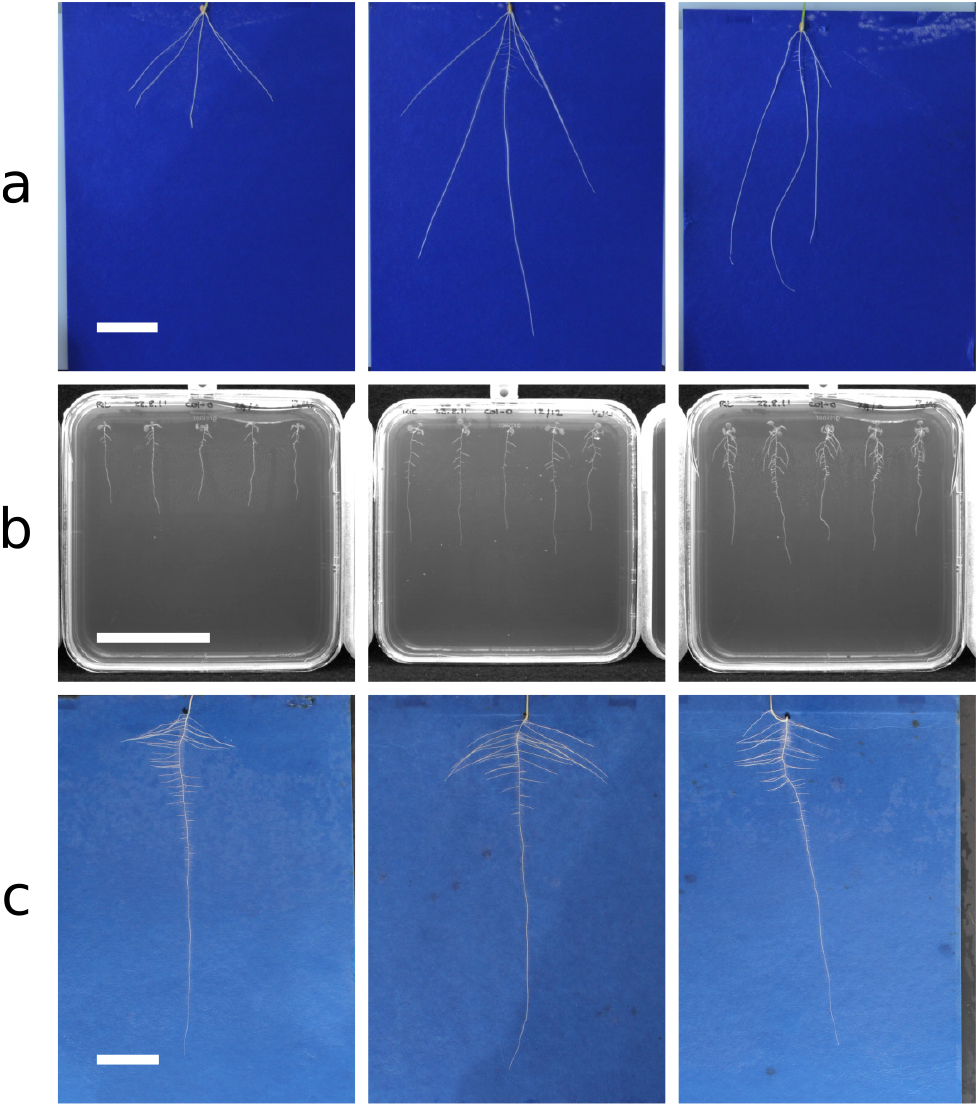
Example images from each of the three datasets used during this work. (a) Wheat (*Triticum aestivum* L.). (b) *Arabidopsis* (*Arabidopsis thaliana*). (c) Rapeseed (*Brassica napus*). Scale bars are 50mm long.

### Transfer Learning Datasets

Our second dataset is composed of images of *Arabidopsis* (*A. thaliana*) grown on agar plates as detailed in [35]. Images of individual plates were acquired using near-infra red imaging utilising the system described in [36]. In this system, multiple seeds are sown on each plate and thus, unlike the primary dataset, each image typically contained up to five plants (Figure 4b). This dataset is considerably smaller, totalling 277 images, and is used as a demonstration of transfer learning with our approach despite limited annotated data. The dataset was split into training and validation sets of 200 and 27 images respectively, and as with the primary dataset, 50 holdout test images were used for final quantitative evaluation.

Our final dataset is composed of images of rapeseed (*Brassica napus*) seedlings, grown in the same system as used in the primary dataset above. This dataset is small, containing only 120 images of individual plants. Our hypothesis is that despite the reduced size, transfer learning from a network trained on both the wheat (similar image background) and *Arabidopsis* (similar root system organisation) datasets will lead to sufficient accuracy. The dataset was split into training and validation sets of 91 and 14 images respectively. We used 15 holdout test images for the final quantitative evaluation. Example images for the two transfer learning datasets can be found in Figure 4b,c.

## Analyses

This section will present a comprehensive performance analysis of RootNav 2.0, including a quantitative evaluation of both the underlying segmentation approach, and the root architecture extraction. We evaluate segmentation accuracy via three common metrics, average pixel classification accuracy (both global and class averages) and mean Intersection over Union (mIoU). We compare segmentation performance of our approach against the well-known benchmark architectures VGG [37], FCN [38], SegNet [39], UNet [40], and DeepLab [41]. We then evaluate the automatic reconstruction of root systems using a comparison of common root phenotypic traits such as the dimensions of the root system, and root counts. For ground truth, we use semi-automatic measurements obtained through expert annotation using RootNav 1.0. Finally, we perform the same experiments to outline the accuracy on the two additional datasets, that contain fewer training images, to demonstrate the efficacy of transfer learning to new species and imaging modalities.

### Root Image Segmentation

RootNav 2.0 is driven by a deep network that segments images of root systems into classes: background, first order roots, and second order roots. Crucial to the accuracy of any subsequent path finding approach is a reliable segmentation. Segmenting whole-root images is important in order to provide sufficient context when distinguishing first or second order roots. Splitting images into efficient tiles reduces memory consumption, but makes distinguishing root type problematic. With this in mind, we designed the network to be efficient by reducing the number of trainable parameters, intermediate feature sizes, and thus overall memory requirements. This allows larger 1024×1024 resolution input. Table 1 shows a comparison of the memory requirements and parameter sizes of commonly used segmentation networks, and our own architecture.

**Table 1.**
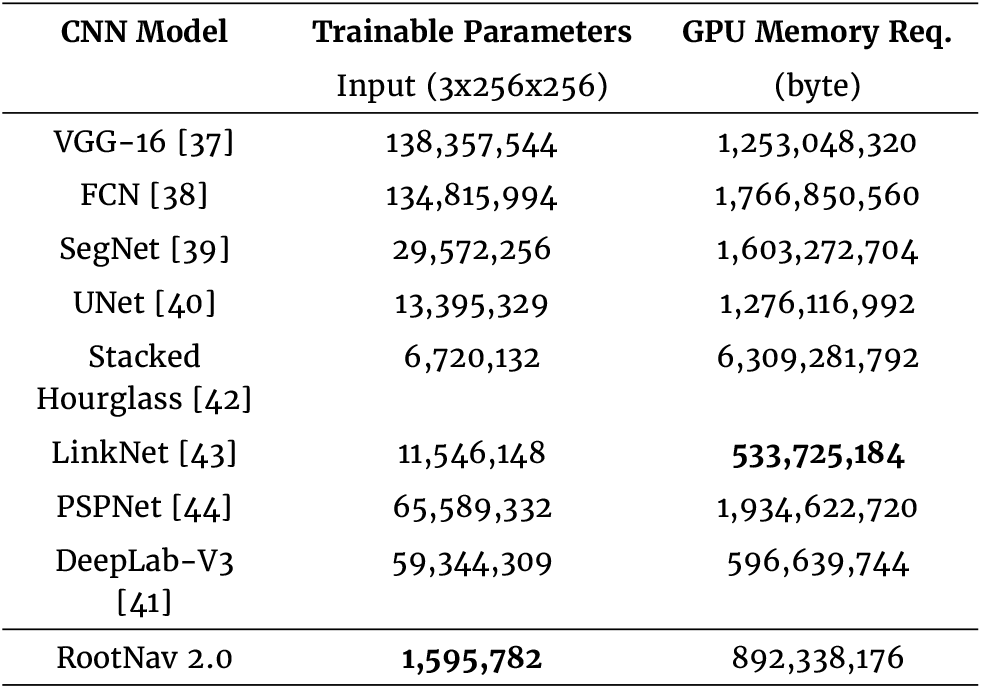
Quantitative Comparison. A quantitative analysis of trainable parameters and memory requirements of different benchmark architectures used during experiments. The input size was set at a constant 3×256×256px size for this comparison.

| CNN Model | Trainable Parameters<br>Input (3x256x256) | GPU Memory Req.<br>(byte) |
| --- | --- | --- |
| VGG-16 [37] | 138,357,544 | 1,253,048,320 |
| FCN [38] | 134,815,994 | 1,766,850,560 |
| SegNet [39] | 29,572,256 | 1,603,272,704 |
| UNet [40] | 13,395,329 | 1,276,116,992 |
| Stacked Hourglass [42] | 6,720,132 | 6,309,281,792 |
| LinkNet [43] | 11,546,148 | 533,725,184 |
| PSPNet [44] | 65,589,332 | 1,934,622,720 |
| DeepLab-V3 [41] | 59,344,309 | 596,639,744 |
| RootNav 2.0 | 1,595,782 | 892,338,176 |

We trained each network on the wheat dataset as described in the Methods. To provide a fair comparison of each network, we allocated 2x Nvidia GPUs with >11Gb onboard memory each for training each network, then trained using consistent hyperparameters such as learning rates, and equal batch sizes. Image resolution was maximised for each network depending on their resource requirements. Accuracy was measured using three standard metrics: Global average accuracy, class average accuracy, and mean Intersection over Union (mIoU). Global average accuracy measures the performance of segmentation over all pixels in the validation set. High values indicate the majority of pixels have been classified correctly. As most pixels are background in root images, high values indicate few false positives, but don’t necessarily demonstrate good root segmentation. Class average accuracy measures the performance of each class separately, before computing a final average. High values here represent good performance across all classes. Finally, mIoU represents the percentage of overlap between each class and the ground truth. Higher values indicate predictions closer to that of the ground truth labelling.

Example image output from each network can be found in Figure 5 with quantitative results for all tested networks across the validation set shown in Table 2. The larger networks contain more features, which while in some cases may improve performance of a deep network, here hinders the ability of each network to resolve finer detail, as they cannot operate at 1 megapixel image resolution. The strong performance of RootNav 2.0 in this experiment may be attributed to its efficient use of features throughout the network, lower memory requirements and thus larger 1 megapixel input sizes.

**Table 2.** A quantitative comparison of the segmentation performance on root images of RootNav 2.0 against other commonly used CNN architectures. Performance is measured using global average accuracy, class average accuracy and mean intersection over union. The classes evaluated are background (no root), first order, and second order roots.

| CNN Arch. | Global avg. | Class avg. | mIoU |
| --- | --- | --- | --- |
| VGG [37] | 37.32 | 38.5 | 31.11 |
| FCN [38] | 43.78 | 48.56 | 39.47 |
| SegNet [39] | 68.68 | 70.23 | 47.05 |
| UNet [40] | 70.47 | 69.65 | 51.88 |
| DeepLab-V3 [41] | 87.7 | 85.90 | 54.0 |
| RootNav.2.0 | **99.6** | **95.1** | **66.1** |

**Figure 5.**
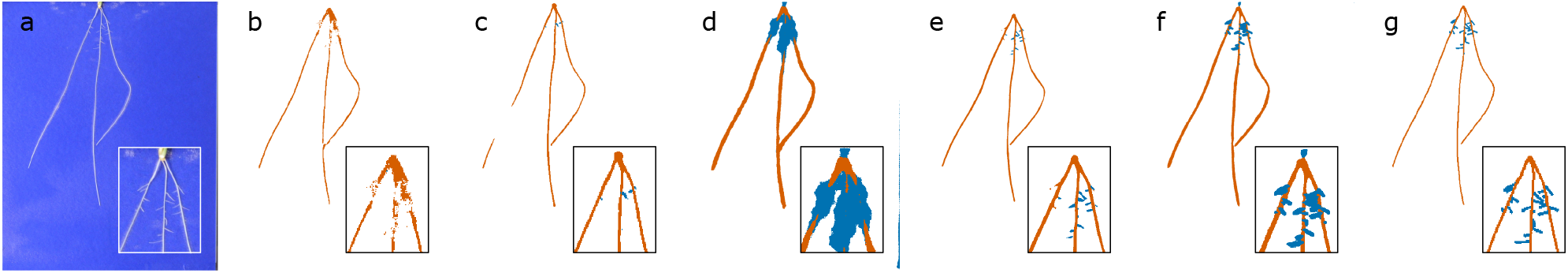
Example image output from each trained network architecture. a) An example hydroponic wheat image. b) VGG [37]. c) FCN [38]. d) SegNet [39]. e) UNet [40]. f) DeepLab-V3 [41]. g) RootNav 2.0

### Extraction of Root System Architecture

After segmentation and feature localization, segmentation masks are converted into a weighted graph structure amenable to traversal with a shortest path algorithm. RootNav 2.0 extracts a full root architecture by performing a series of heuristic searches across the image. First, shortest paths are found between all first order root tips and the most appropriate seed location (defined as the seed first reached during a heuristic search). This generates a series of first order roots, to which second order root paths are found from all second order root tips. The output of this process is a complete root architecture description, stored in RSML format, from which phenotypic traits may be derived. We compare the output of RootNav 2.0 against ground truth measurements captured using RootNav 1.0 in collaboration with an expert user. Quantitative traits were measured directly using the RSML output by both tools; results may be found in Figure 6.

**Figure 6.**
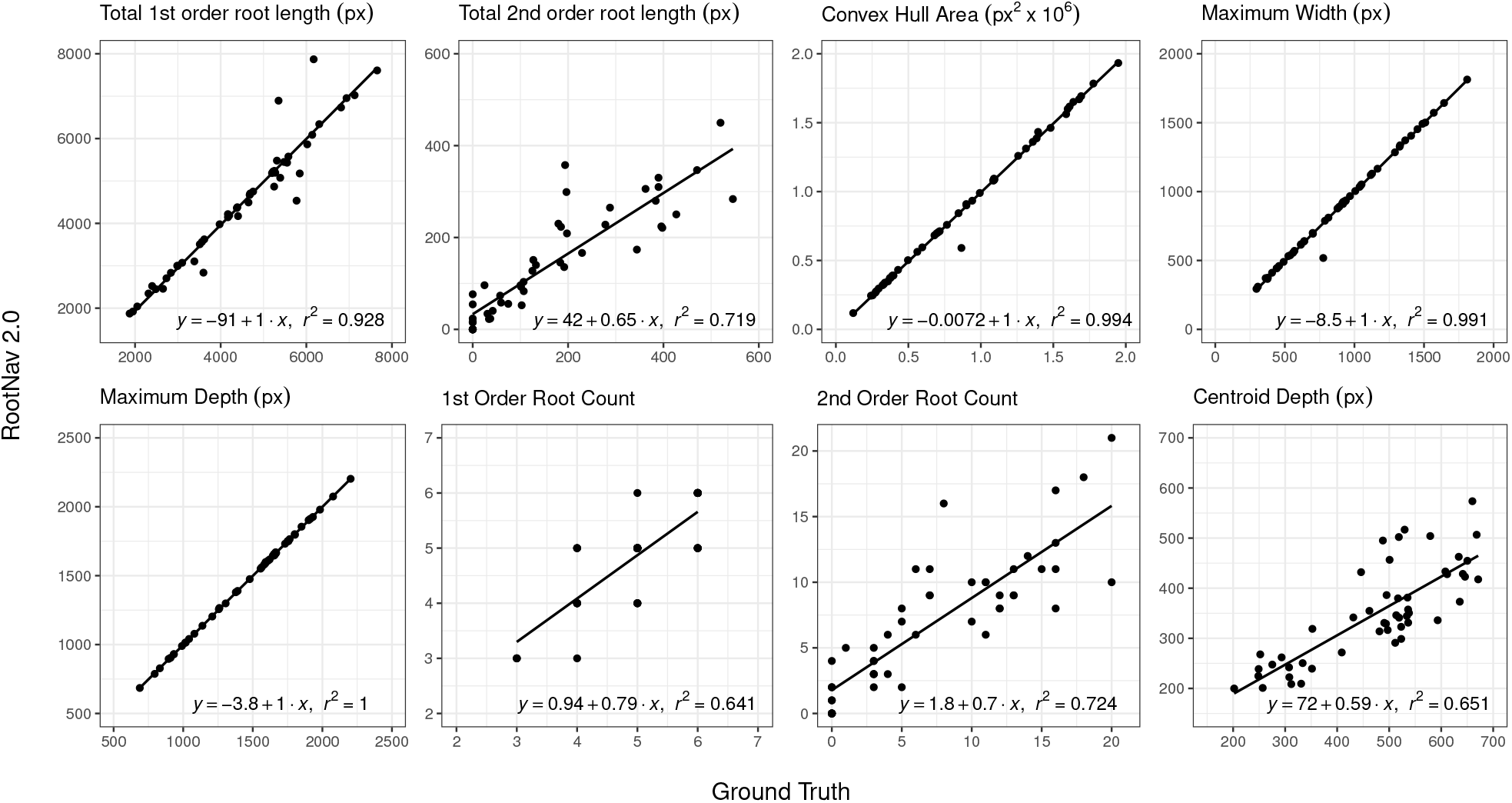
Numerical results showing a range of root system traits measured in RootNav 2.0 against ground truth measurements on the wheat test set. For each trait we also fit a linear regression model and report the *R*^2^ value.

We chose a range of root traits that are both representative of the measurements commonly used in the root phenotyping literature, but also ones that exercise various aspects of our particular approach. For example, we include traits that measure the accuracy of feature detection (e.g. total root count) and those that also measure the accuracy of the shortest path approach (e.g. total root length). In Figure 6 it can be seen that there is strong agreement between the results of RootNav 2.0 and the ground truth measurements. Measurements based on the extremities of the root system (maximum depth, maximum width, and convex hull area), produced values very close to those in the ground truth, with *r*^2^ values above 0.99. Traits that summarise the entire root system, such as centroid depth, provided *r*^2^ values in the range 0.64-0.72.

The prediction of first and second order root counts achieved *r*^2^ values of 0.641 and 0.724 respectively. For first order roots, we observed that the majority of incorrect predictions were either one count higher, or one count lower than the ground truth, and that these confusions often occurred near the seed position, where a seminal root may visually appear similar to a second order root that emerges near the seed, or vice versa. Other failures were produced by roots leaving the field of view of the camera, but that had been annotated by the expert, or where two root tips grew in very close proximity (within a few pixels). Second order roots were typically much shorter, and often in close proximity. Some missed root tips would be caused by non-maximal suppression, when the R-Tree data structure is used to remove possible duplicates. We also found that the contrast on second order roots was lower, as they were usually thinner, which might account for some missed tips in this class. Errors in the detection of root tips will also propagate errors into the total root length measurements, since these roots will not be detected. For second order roots, we found that most of the error in root length may be attributed to missed roots, rather than errors in path finding. For primary roots, path finding was usually robust, except in cases where two roots grow side-by-side. RootNav 1.0 handled these errors by allowing a user to intervene and correct any mistakes, in RootNav 2.0 we wish the process to remain fully automatic, and so we do not explicitly correct for this. However, the occurence of this type of growth is in the minority, in our experience. Centroid depth is measured as the mean position of all roots, and so is influenced by the detection and path finding of every root.

An understanding of where and how RootNav 2.0 may produce errors provides insight into these results. An accurate measurement of maximum depth depends on only two variables, the location of the seed, and the location of the first order root tip, that is lowest (in terms of y-position) in the image. The graph of maximum depth in Figure 6 reflects the fact that these two features are successfully found in every case. Similarly, maximum width depends only on the left- and right-most roots, and convex hull only on the outermost roots throughout the architecture. A missed second order root within a root system will not affect these traits, so these results are robust even where some roots have been missed. This tells us that for the majority of images, the locations of the seeds, lowest tips, and outermost roots are detected successfully, and that these traits that measure the extremities of the root system are robust.

### Transfer Learning to New Species and Images

To demonstrate the adaptability of our approach to different species and imaging modalities, we retrained the network first on an *Arabidopsis* dataset, comprising approximately 277 images of *Arabidopsis thaliana* grown on agar plates. We then trained once more from the wheat dataset to the rapeseed dataset, comprising 120 images of *Brassica napus* on germination paper. In both cases we extracted RSML root descriptions, and quantified these in the same way as the wheat dataset. We also trained both networks from randomly initialised weights, rather than transfer learning, and found that the datasets were too small to train effectively (Supplementary Figures 6 and 7).

#### Arabidopsis thaliana

The *Arabidopsis* dataset contains marked differences from the wheat data. The dataset contains many fewer images, which makes transfer learning essential to avoid overfitting. This species has a taproot structure that contains a single primary root, from which lateral roots emerge, rather than multiple first order roots in the form of a primary and multiple seminal roots. This dataset is also imaged on under infra-red illumination and so contains no colour information, and a very different background arrangement consisting of a plastic plate containing semi-transparent growth media instead of blue germination paper. Finally, each plate typically contains five plants, rather than a single plant. We found that the network and heuristic searches adapted well to this new domain. Quantitative results are shown in Figure 7, with full results found in Supplementary Figure 1, and example image output in Supplementary Figure 4.

**Figure 7.**
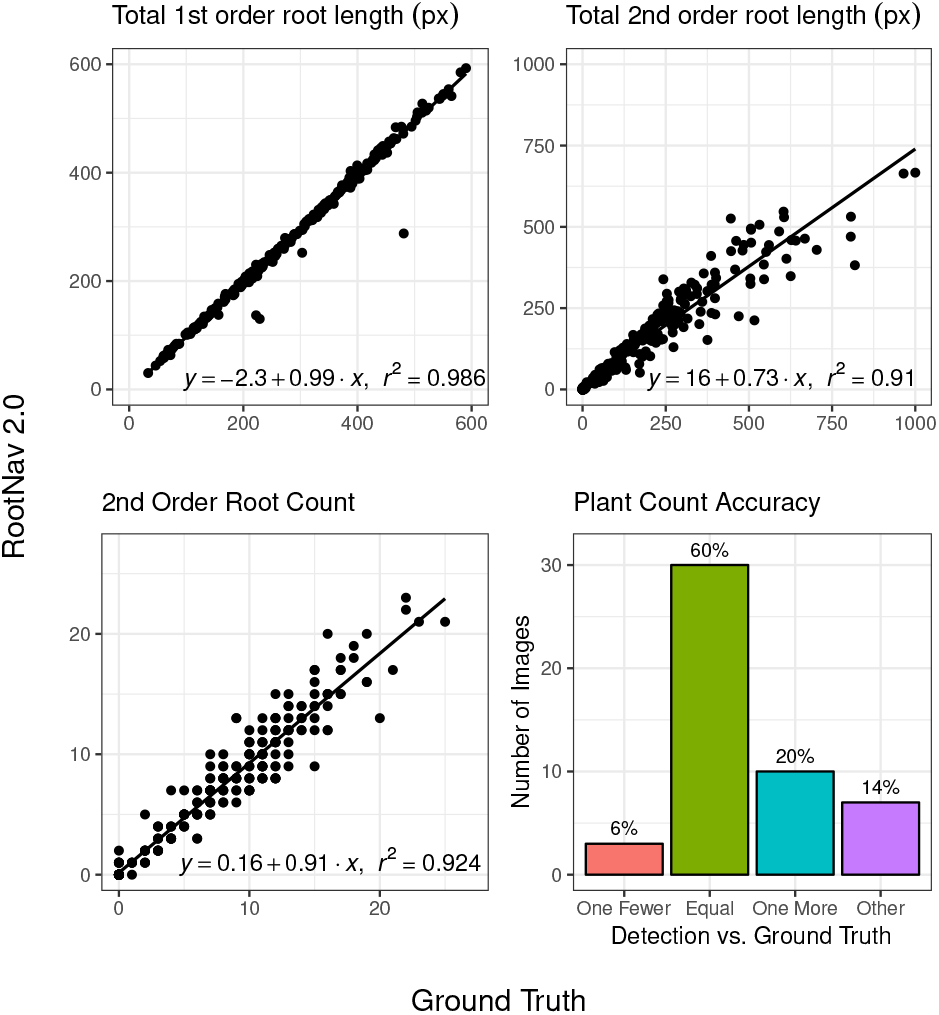
Numerical results showing a sample of root system traits measured in RootNav 2.0 against ground truth measurements on the ***Arabidopsis*** test set. Each image contains up to 5 plants, and results are presented per-plant. For each trait we also fit a linear regression model and report the *R*^2^ value.

Despite the smaller number of images available for training, the results show a good performance after transfer learning to the new data. Not every plant was successfully detected; a missing or additional primary root tip or seed location would mean that the number of plants was under or overestimated. In 60% of images examined, the tool correctly identified the same number of plants as were marked in the ground truth. In 6% of the images, a single plant was missed, usually due to the plant being extremely underdeveloped, but having been annotated by the user anyway. We found only a single instance in one image that contained a well established plant which had not been identified by our network. Over-counting of plants was more common, with 20% of images identifying an additional plant, and 14% identifying more beyond this. In the majority of cases we found that these errors were caused by unusual angles in the leaves and germinated seeds at the top of the plant, producing false positives. This is something that would likely be corrected with additional training data; remember, we are using a very small amount of training data for this image class, versus the wheat images. Where duplicate plants were found, they were often extremely close to, or even above, an existing plant location. These duplicates could be removed easily via post-processing; this is something we do not address in this paper.

Of the plants that were successfully identified, the traits captured by the tool offer a good agreement with the ground truth. As with the wheat dataset, measures of the extremities of the root system such as maximum depth performed with the highest accuracy, but we also found that total 1st order root length was very close to the ground truth in the majority of cases. Errors here usually indicated a second primary root incorrectly detected alongside an existing one, a feature we do not yet remove in post-processing as with duplicate plants, though this would be possible. The detection of second order roots was also highly correlated with the ground truth measurements, and the total length of all second order roots (measured per plant) correlated with the ground truth with an *r*^2^ of 0.91.

#### Brassica napus

This dataset uses the imaging format of blue germination paper with single plants (like the wheat dataset), but contains the same species root structure as *Arabidopsis* (a single tap root from which all other roots derive). This dataset contains the fewest images, with only 90 images used for training. We use this small dataset as a demonstration of the efficacy of transfer learning, but we also note that training over a slightly larger dataset in practice would be worthwhile for improving robustness. Results may be found in Figure 8, and in full in Supplementary Figure 2.

**Figure 8.**
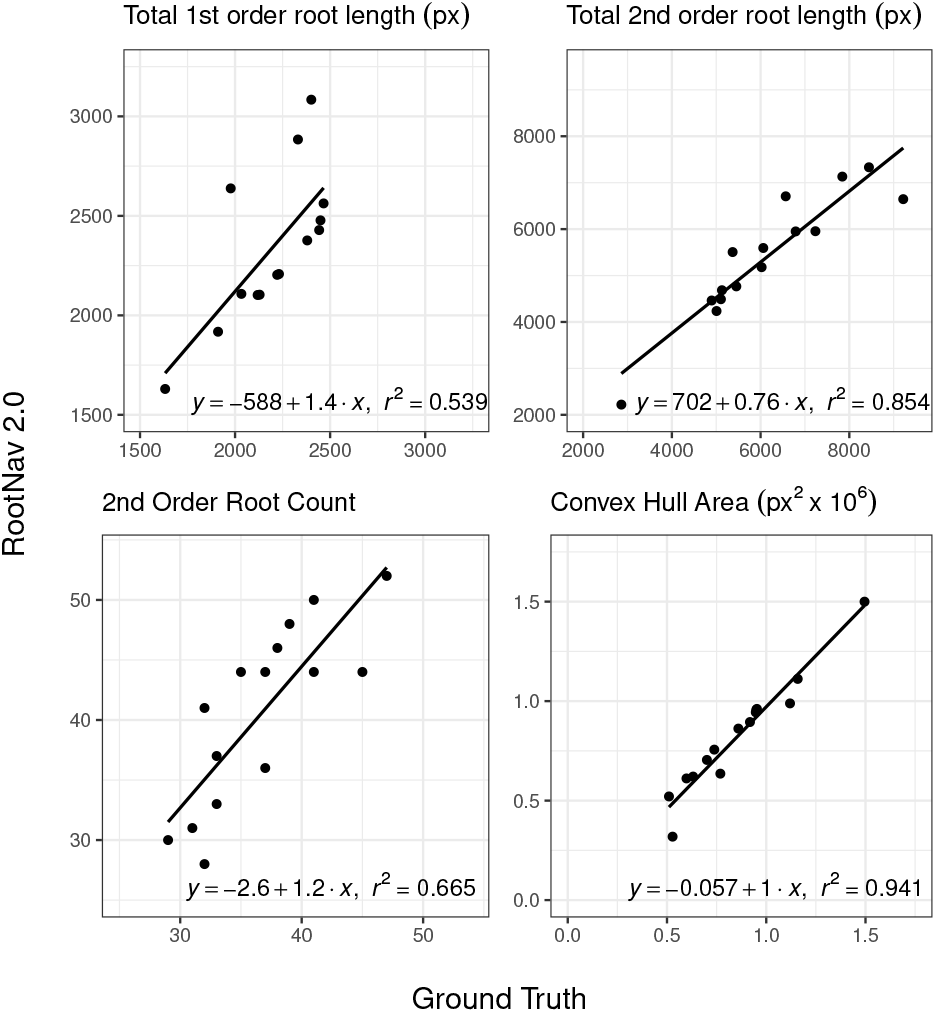
Numerical results showing a sample of root system traits measured in RootNav 2.0 against ground truth measurements on the ***Brassica napus*** test set. For each trait we also fit a linear regression model and report the *R*^2^ value.

It can be seen that the correlation between RootNav 2.0 and ground truth ranges from *r*^2^ of 0.539 (first order root length) to 0.941 (convex hull area). As this is such a small dataset, the test set contains only 15 images, meaning that there is inevitably more noise in the results than the previous experiments. Nevertheless, the results are promising, particularly given the tiny size of this dataset compared with typical standards for deep learning. As with both previous datasets, convex hull and other extremity-based measures provided the most reliable results. Accuracy of the total length and root count metrics had a lower *r*^2^ than the other datasets, caused we believe by the smaller training set meaning that the approach is a slightly less robust to noise. We found that in a few images the longest lateral tips were incorrectly classified as first order, causing erroneously-high measures of first order root length. We believe this occurred where these laterals are mistaken for wheat seminal roots, on which the network was originally trained. This occurred on the minority of test images, and is a problem we are confident would be resolved with a more training data. It would be possible to use prior domain knowledge, for example the knowledge that rapeseed has a single tap root, in order to clean the output during post-processing; as with the *Arabidopsis* dataset, we do not perform any post-processing of this kind in this work.

After segmentation, the structure of these root systems were quite amenable to traversal using a shortest path approach. In many cases the longest roots grow close together, which causes errors where a search may travel along the same path as another root. We found this did not substantially increase the error in total root length, as many of these roots grew in close proximity, and were of similar length. Nevertheless, dealing with root overlap in an efficient and automatic way is a topic worth exploring in future work.

### Performance Analyses

We measured the time taken for both tools to complete the full pipeline, from image to RSML output. We timed RootNav 1.0 by annotating random images from each test set from scratch; the total time taken to annotate 10 images of wheat and *Arabidopsis*, and 5 rapeseed images were recorded, and averages computed. The annotation was performed by an expert user who has many years of familiarity with the tool. For RootNav 2.0, we processed each test set and then calculated the average inference time per image. Results may be found in table 3.

**Table 3.** A performance comparison of RootNav 1.0 against RootNav 2.0. The time to process a random sample of images from each test set was measured, and an average time per image calculated.

| Dataset | Average Processing Time (s) |  |
| --- | --- | --- |
|  | RootNav 1.0 | RootNav 2.0 |
| Wheat | 68.8 | 8.2 |
| Arabidopsis | 109.8 | 7.5 |
| Brassica | 132.4 | 14.0 |

In both systems a more complex root architecture typically leads to a longer analysis time. In RootNav 1.0 this is due to the human input required, with RootNav 2.0 the path finding takes longer if there are more lateral roots, or roots are longer. Regardless, for each data set RootNav 2.0 offers a substantial speed advantage over the original tool. It should also be noted that the time presented here for RootNav 1.0 requires that the user engage with the software continuously. Since RootNav 2.0 is fully automatic, the *human time* cost is essentially zero, as images could be batch processed overnight. This test was also run on a single CPU and GPU, where additional computational resource would linearly scale the speed of the system. If performance was a serious consideration, a dedicated parallel hardware setup could streamline RootNav 2.0 performance considerably.

## Discussion

In this paper we have introduced RootNav 2.0, a state-of-the-art, fully automated root phenotyping tool. It is powered by a deep convolutional neural network in an encoder-decoder configuration, designed to perform segmentation efficiently in high-resolution images. The network segments root from background, and can distinguish first and second order roots. This deep learned root segmentation provides a strong foundation upon which users may derive common architectural traits, such as those based on RSA skeletonisation. We have adapted the network, however, to simultaneously predict the location of key root architectural features: the seed location, and first and second order root tips. This knowledge then drives a heuristic search that reconstructs the entire root system. This topology is represented as spline curves, and output in RSML format.

A quantitative analysis of RootNav 2.0 shows it offers comparable accuracy against the original RootNav on large training sets. Over a range of standard trait measurements the new tool produced highly correlated results against the ground truth, with *r*^2^ values ranging from 0.64 to 1. Performance on traits representing the bounds of the root system were among the highest *r*^2^ values. On smaller datasets, we have demonstrated that transfer learning produces accurate results despite many fewer training examples. This adaptability is a key advantage for those within the research community looking to use RootNav 2.0; those that use different growth conditions, image capture approaches, or require the analysis of different species may adapt one of our existing trained models with a minimum of effort using transfer learning.

RootNav 2.0 is substantially more convenient to use than previous semi-automated tools, with no human interaction required at any point during the pipeline. The entire process requires approximately 15 seconds processing time per image. The output may be analysed using the RootNav viewer tool, or any compatible RSML analysis pipeline. Training the original network took a few days on suitable hardware, with transfer learning to a new dataset typically taking about half a day. All code, trained networks, and detailed documentation on use and retraining will be made available at the links below.

We believe that RootNav 2.0 will prove to be a key milestone in root phenotyping, further encouraging the uptake of machine learning in addressing these important challenges. In future work, we will continue to adapt this approach to new and varied datasets, maximising the potential for use in the research community. We will also explore the use of more robust heuristic searches, combined with appropriate segmentation output from the network, to address the challenge of crossing and intersecting root systems. We will also develop approaches to ease the sharing of network models, and indeed the retraining process required to adapt them to specific scenarios.

### Potential implications

We believe that RootNav 2.0 offers a substantial increase in accuracy over bottom-up approaches to root image analysis. It also offers an increase in throughput over existing semi-automatic tools. Importantly, results with the *Arabidopisis* dataset suggest the approach will be applicable to images obtained with other phenotyping systems such as rhizotrons. With continued community support, RootNav 2.0 has the potential to be the first true species - and platform - agnostic analysis tool in the plant sciences. This will provide researchers with the ability to analyse root systems at larger scales than ever before, at a time when large scale genomic studies have made this more important than ever.

## Availability of source code and requirements

Project name: RootNav 2.0 Project home page:

https://github.com/robail-yasrab/RootNav-2.0

Operating system(s): Platform independent Programming language: Python, C#

Other requirements: Python 2.7, PyTorch 1.0.1, License: GNU GPL

Any restrictions to use by non-academics: None

## Availability of data and materials

Datasets will be made available at

https://plantimages.nottingham.ac.uk upon publication.

## Methods

### Training, validation and test image preparation

For each image we obtained ground truth annotations using the original RootNav 1.0 software. This software is semi-automatic, and allows users to manually intervene to correct errors in either segmentation or RSA extraction. We are using this data as a ground truth, rather than to evaluate the accuracy of RootNav 1.0, and as such annotators were instructed to spend sufficient time on each image to correct all mistakes they could identify. This semi-automatic process often requires a large amount of human interaction, and is time consuming, but the approach has provided very reliable ground truth annotations. All ground truth was stored in RSML format.

RSML data for each image was converted into a series of segmentation masks and feature heatmaps for use in training. Segmentation masks were created separately for both first and second order roots by rendering them as polylines over a blank image. RootNav 1.0 does not measure diameter information for root systems, but the seedlines are sufficiently young that root diameter is quite consistent across species and images. We rendered each root with a width of 8 pixels. For heatmap output, the seed location, first and second order root tip locations were rendered as in [5], as separate images of blurred Gaussian points of standard deviation 1.0 pixels. The result of these processes is that for each input image there are five associated output images, two segmentation masks for first and second order roots, and three heatmaps for seed position, first and second order root tips Fig. 9.

**Figure 9.**
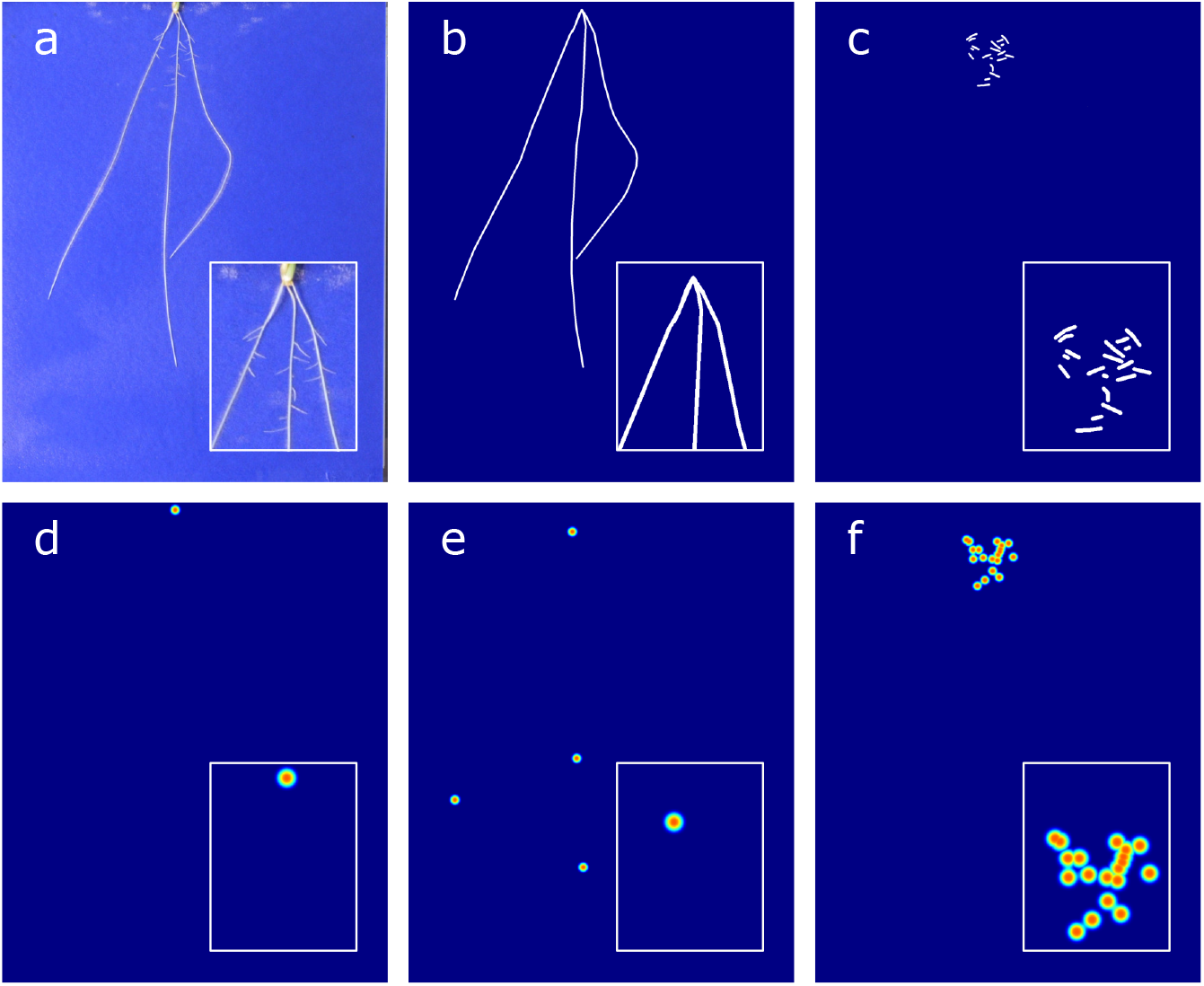
Sample training image and ground truth, zoomed sections and colour added for clarity. a) An example wheat image. b) Segmentation mask for first order roots. c) Segmentation mask for second order roots. d-f) Heatmaps for seed, first, and second order tip locations.

At this point, we have constructed three suitable training sets of images based on manual annotations. The next task is to construct a suitable encoder-decoder architecture capable of segmenting these images and locating root features.

### CNN Design

#### Input and Output Resolution

We used the PyTorch [45] framework to develop the network, training and validation code that drives our segmentation approach. The network is based around an encoder-decoder architecture (Fig. 2), but has been adapted to handle the higher-resolution images seen in the datasets. Encoder-decoder CNNs are memory intensive, particularly at points towards the start and end of the network where the spatial resolution is high. Each layer calculates many features, which each exist as an image stored in memory. Over many layers, the computational cost becomes prohibitive. Previous work, such as [5] used small input and output sizes of 256×256 pixels. Other commonly used networks such as VGG-FCN [38] and U-Net [40] use similar input sizes. Root images pose a challenge in this situation as roots may be only a few pixels in diameter, but exist as part of a large, connected architecture covering many megapixels. Shrinking the image into a convenient size will make processing simpler, but also badly degrade the quality of these small features. In scenarios such as this, where shrinking the input this far may represent a significant loss in quality, it is common to tile the input into small cropped sections, and run the network repeatedly. This is the approach taken by [5], in which wheat images are tiled, processed, and then reconstructed. The drawback of tiling images is that each tile is then considered in isolation, removing vital context on its position in the wider image. In the root datasets, for example, first and second order roots often appear identical when not viewed as part of a larger architecture.

In this work, we limit the size of input images to 1024×1024 pixels. For the wheat and rapeseed datasets, this requires down-sampling of the input and output images, but only by a moderate amount, in which fine root detail is preserved. For the *Arabidopsis* plate images, no downsampling is required as they are already of a suitable size. Upon completion of the deep learning, images are returned to native resolution to ensure that the output measurement scale is preserved.

#### Network Architecture

Our complete network operates on input images of 1024×1024, and outputs segmentation masks and regression maps of 512×512. A diagrammatic overview of the network may be found in Figure. 10, with a description of the layers in Table. 4. The core of the network is an hourglass architecture similar to those used in [42, 5], but here we use a restricted number of features throughout, and do not use stacked structure (repeated encoder-decoders after one another). These alterations to the network allow it to successfully process the 1 megapixel input size without reaching the limit of available memory. We also perform additional downsampling and upsampling at either end of the network. Initial strided convolutional layers with large filter sizes of 7×7 are used to extract features and downsample the image size, before interleaved residual blocks and max pooling operations are used to further reduce the spatial size of the input to 128×128 pixels. The hourglass architecture performs the primary encoder-decoder role, with downsampling performed using max pooling, and upsampling performed using bilinear interpolation. The output of the hourglass is a set of 128×128 pixel feature maps, after which learned deconvolutional filters and residual blocks are used to return to a 512×512 pixel spatial resolution. Finally, two paths are used to separately predict segmentation masks and feature heatmaps. Each branch comprises 1×1 convolutional layers for prediction, with the segmentation output also passed through a sigmoid output, as required by the binary cross entropy loss function.

**Table 4.**
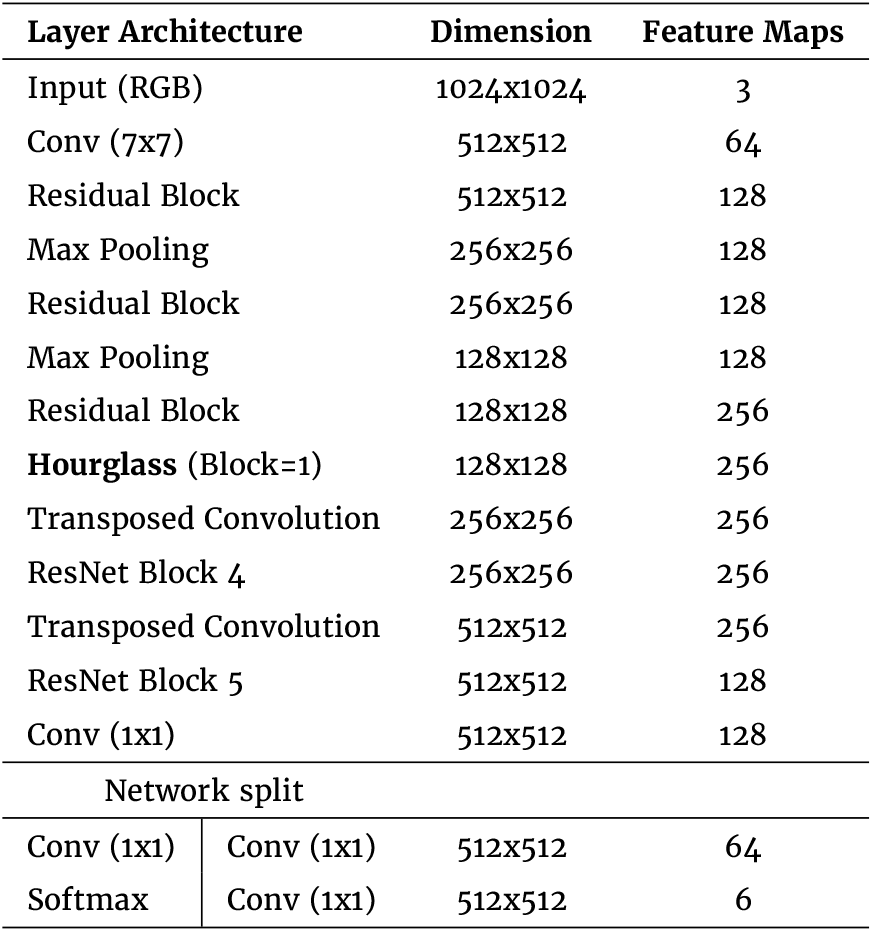
The proposed CNN’s Layers. Batch normalisation layers and ReLU activation functions are used between layers and within residual blocks. The hourglass used is equivalent to a single stack of the type used in [5].

**Figure 10.**
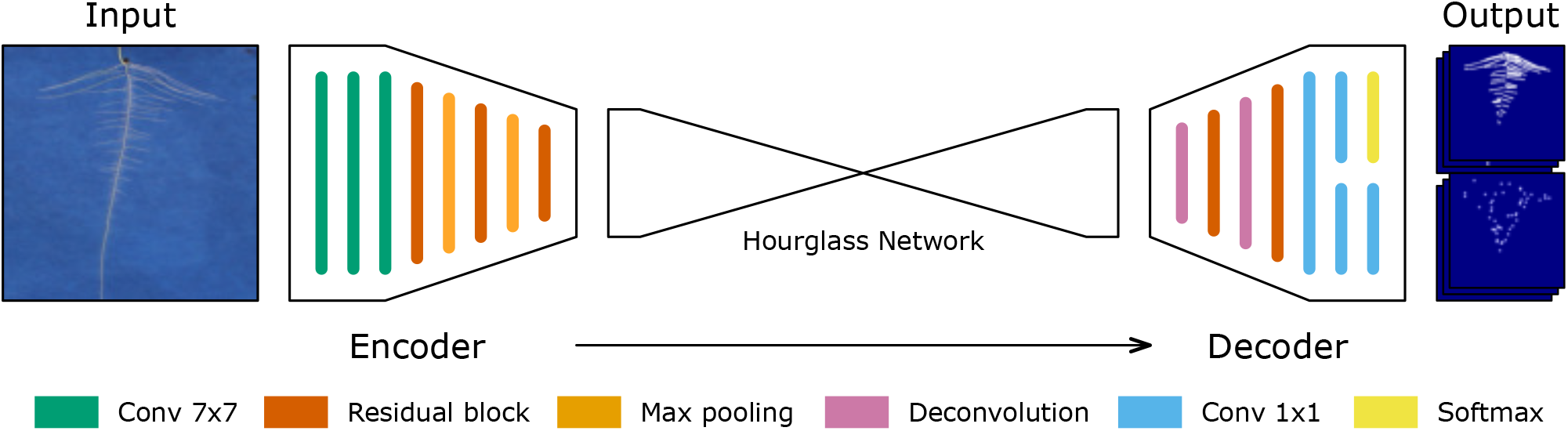
The network used is an extended encoder-decoder architecture. The input is filtered and downsampled into an efficient size, before an hourglass network [42] performs the remaining encoding and some decoding. Finally, a series of learned deconvolutional layers upsample back to 50% of the original input size. The network is split into two fully-convolutional branches at the output, which are responsible for learning the segmentation and heatmap regression outputs separately.

### Loss Functions

The output of our network is divided into two paths with different objectives. The first outputs segmentation masks containing locations for first and second order roots. Each of these is a 2D binary output, and is trained using a cross entropy loss. It is common in root images that the number of background pixels heavily outweigh the foreground. Calculating a loss over an unbalanced dataset such as this is likely to cause a bias towards background pixels, causing error and underseg-mentation of the foreground. We apply a class balancing approach to the standard cross entropy loss, based on median frequency balancing [46]. Weights are assigned to each class inversely proportional to the median frequency in which that class appears throughout the entire training set. This reduces the weight for classes that appear more often, in this case background, and increases the weight of foreground classes such as first order roots. The accuracy of classes with higher weights are prioritised during training. The proposed loss *L*_1_ is given in Eq. 1.

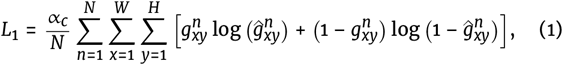

where for *N* features, 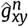 is the predicted class output at location (*x*, *y*), and 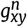 is the ground truth prediction at that location. The weight of each class is scaled by it’s frequency relative to the median frequency of all classes by α_*c*_, given by:

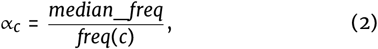

where *freq*(*c*) is the frequency of occurrences of pixels of class *c* divided by the number of pixels in any image containing that class, and *median_freq* is the median of these frequencies over all classes.

The second path is responsible for predicting key feature locations on the root system. Specifically, the seed location, first order root tips, and second order root tips. The output is three 2D outputs, trained using a mean squared error loss, predicting likely locations for root features, represented by 2D Gaussians centred at each feature location. The proposed loss *L*_2_ is depicted in Eq. 3.

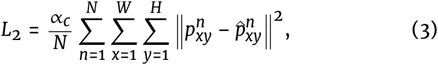

where for each of *N* features, 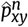 is the predicted feature likelihood at pixel (*x*, *y*), and 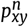 is the expected ground truth at the same location.

The final loss *L* = *L*_1_ + *L*_2_ trains the network end-to-end, balancing the objectives of both paths. We found that additional scaling factors applied to the loss of either path was not necessary for accurate training.

### Training

Beginning with the wheat dataset, the network was trained end-to-end from scratch using the rmsprop opimizer. The initial learning rate was set to 1*e*^−4^ and reduced by a factor of 10 after 50,000 iterations. The network was trained using a batch size of 6 for 500k iterations, although we found performance plateaued after approximately 400-450k iterations. During training, we selected the best performing model from the validation set.

We applied random augmentation to the training set in order to reduce potential overfitting. We added random horizontal flipping with a 50% probability during training, as well as random rotation of in the range [-30°,30°]. We experimented with random cropping as in [5] but found that cropping often caused the removal of parts of the root system, sacrificing context crucial, for example, in distinguishing first and second order roots. For this reason we did not use random cropping during these experiments.

#### Transfer Learning

Transfer learning is the process of training on new data by beginning with an existing trained network’s parameters, rather than randomly initialised weights. We began by training the wheat network to completion. This is a large dataset, with more than sufficient images to train a network reliably from scratch. As noted in the above section, successful training simply means acceptable performance on the validation images. We began with the existing wheat network and retrained on the *Arabidopsis thaliana* dataset. This is a smaller dataset, but the use of pre-trained weights allows a network to make use of any useful image filters learned during the initial training. We experimented with training from scratch on the smaller dataset, but found that we were unable to train a network that performed reliably on the validation set (Supplementary Figures 6 and 7). The same process of training and validation was used to complete training on the new dataset, except that we limited training duration to 120k iterations.

Finally, we repeated transfer learning from the wheat dataset to the *Brassica napus* data, which has the fewest images of the datasets we use. We explored training from scratch, as well as using pre-trained weights from either the wheat or arabidopsis datasets, and found the wheat network offered the most reliable starting point, a fact we attribute to the similar background and foreground colours, and scales for both datasets. As above, we trained for 120k iterations and selected the model with highest validation performance.

### Post Processing

#### Dense CRF

Each segmentation mask was passed to a Dense Conditional Random Field (CRF) to improve smoothness and maximise agreement between similar neighbouring pixels. We found this approach has a subtle but helpful effect on the separation between roots growing in close proximity, and the smoothness of the boundaries of segmented roots. We used the dense CRF proposed in [47], in which each pair of random variables (pixels) are connected by an edge [48], weighted using a Gaussian pairwise potential. The effect was a smoothing of conflicting regions of pixels where the image was cluttered; but where segmentation was already successful, the approach had no notable negative impact on the results.

#### Feature Localisation and Non-maximal Suppression

The heatmap regression output contains the probable locations of the seed, as well as first and second-order root tips. These are represented as 2D Gaussian distributions, the centre of which lies on likely feature locations. We obtain a discrete location for each feature via a non-maximal suppression (NMS) approach [49], which suppresses all predicted pixels except those that are greater than their surrounding neighbours. For a Gaussian-based distribution, this has the effect of locating the centre of the distribution. Our implementation of NMS uses pixel-wise search, immediately discounting any pixel output below a pre-defined threshold (in our case, 0.7), for speed. Each pixel is then compared with its neighbor pixels in a 3×3 window, where the central pixel “c” is non-maximal, if another pixel of greater or equal intensity is discovered in its neighbourhood, the algorithm skips to the next pixel in the scan line [50].

For the majority of root tips in isolation, NMS will successfully return a single location for each true root tip. This relies on the heatmap regression layers of the network returning well-formed Gaussian dsitributions in all instances, which while likely, may not occur in the presence of image clutter, confusing root hairs, or multiple tips in close proximity. To avoid two positions being returned for a single underlying root feature, we identify and suppress neighbouring features. We use an R-Tree data structure to efficiently query for neighbours within close proximity. When NMS returns a new position on the image, the R-Tree is searched for nearby features that have already been added, and prevents locations from being added twice. In our experiments, we considered a new position a duplicate if it falls within 8 pixels radius of an existing feature, which is derived from the scale of the roots in our datasets.

### Root Architecture Reconstruction

After successful pixel-wise segmentation of the complete root system architecture and extraction of the tips and seed locations, we are now able to reconstruct the whole root skeleton. This procedure is similar to the original RootNav 1.0 tool [9], except it is now driven by more accurate class-aware segmentation, rather than error-prone root likelihood estimations. RootNav 2.0 can place more reliance on the accuracy of the segmentation, and make use of each segmentation map separately to ensure that roots are not traversed over the wrong material, e.g. that first order roots prioritise image locations of that class. We establish an 8-way connected graph structure throughout the image, where the weights travelling to neighbouring pixels are calculated as a function of their class, the path we are trying to find, and the distance between them. Each segmentation mask is converted into a distance map of values [0, 1] indicating the distance from any background pixel. We then convert this distance into a weighting that prioritises paths along root centres; the maximum weight we assign to any root pixel is 0.05, for pixels near the root edge. The weight decreases towards the centre of the root, to a minimum of 0.01. Since the graph includes diagonal connections, these are weighted by an additional cost of 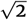 to account for the longer distance. Finally, any pixel that does not belong to the specific class being traversed, e.g. first order root only, is assigned a weight of 10.0, representing a much stonger penalty for traversing these pixels. Unlike RootNav 1.0, we use separate graphs and searches for first and second order roots. A value of 10.0 was chosen simply as a very large increase in weight when compared to the minimum cost for any segmented root material. Different weight values are effective, as long as they are large enough relative to root material to avoid the shortest path taking shortcuts across background pixels where this is unnecessary. In practice, these weights are only traversed if there is a gap in the segmentation for true root material.

A* search [51] is a path finding algorithm that in our implementation seeks to find a path of minimal cost between locations on a root system. It is an extension of Dijkstra’s shortest path algorithm [52], and along with distance travelled also considers a heuristic measure of the remaining distance to the goal. Pixels are explored based on the lowest cost first, in order to minimise the function

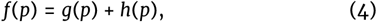

where *g*(*p*) is the sum of all weights to *p*, and *h*(*p*) is the remaining distance, which we calculate as the Manhattan distance, or *L*^1^-norm.

In the case of RSA traversal, minimal cost paths between key features such first order root tips and seeds represent reconstructed roots. A* searches are initialised from all first order root tips, travelling along segmented roots until they reach any seed point. Upon reaching a seed location, the entire path is recorded as a first order root. Once all first order roots have been traversed, a new series of searches are begun from second order root tip locations, ending at any encountered first order root. In RootNav 2.0, we assume that the closest first order root material connected to a second order root tip is the emergence point of that root. The output of each search is a list of pixel co-ordinates representing the individual roots within the RSA.

#### Spline fitting

The use of a distance map that prioritises the centre lines of roots generally acts to smooth the paths found throughout the root system. This may not be the case where there is noise in the segmentation output, or roots cross, and the distance map is less reliable. We smooth each root path using a spline curve representation. Control points are sampled at equal spacing along each path, before the path is re-sampled using cubic splines. Each spline includes a tension parameter that we set at a constant of 0.5 for our experiments. Both the spline and a polyline representation are output into the final RSML file to ensure maximum compatibility with other tools.

#### RSML and Output

The RSA reconstruction approach in RootNav 2.0 does not perform phenotypic measurements itself, rather it extracts a root topology along with segmented images from which traits may be derived. The entire root system for each plant in an image is exported using the Root System Mark-up Language (RSML) format [33], providing a standard and interoperable format. RSML is an XML document specifically designed to store 2D and 3D root architectures. It also stores meta-data, plant properties, and is compatible with numerous analysis tools. Some existing plant phenotyping tools offer RSML import support, meaning that they may also load root systems created automatically using RootNav 2.0. Our approach also outputs first and second order root segmentation images, representing an alternative source of quantitative data. Many tools such as Ez-Rhizo operate on such images, but the segmentation masks generated here contain very little image noise, making them more amenable to further automated analysis. An example output may be seen in Figure 11. For this work we performed quantification entirely using the RSML output. Phenotypic measurements were calculated from each RSML file using the existing RootNav Viewer tool, which has been extended and updated for this publication.

**Figure 11.**
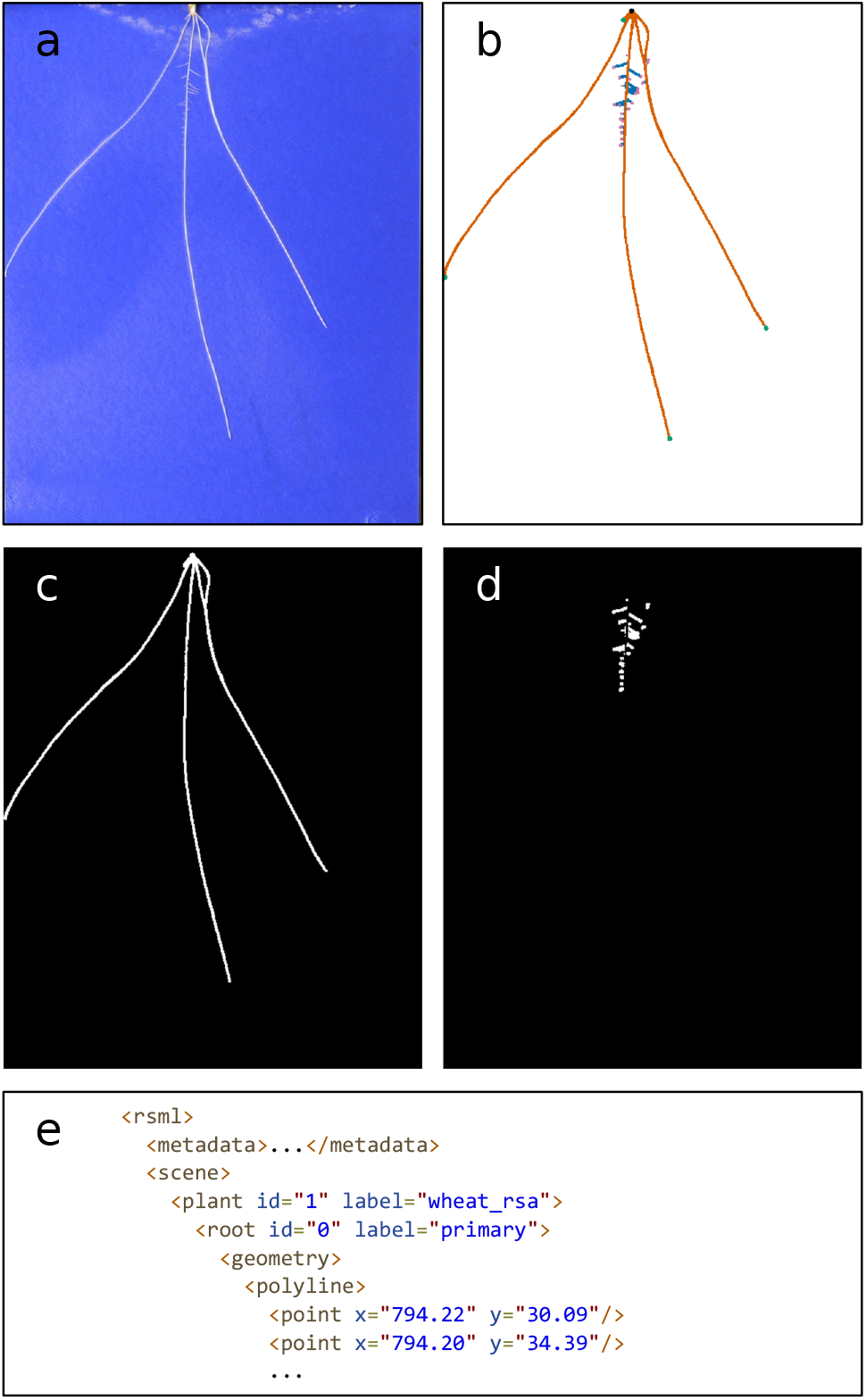
Example output from *RootNav 2.0*. (a) Input image. (b) Colour-coded segmentation mask. (C,d) Binary segmentation masks for first and second order roots. (e) A sample of the RSML file representing the entire architecture.

