## Supplementary Figures for "RootNav 2.0: Deep Learning for Automatic Navigation of Complex Plant Root Architectures"

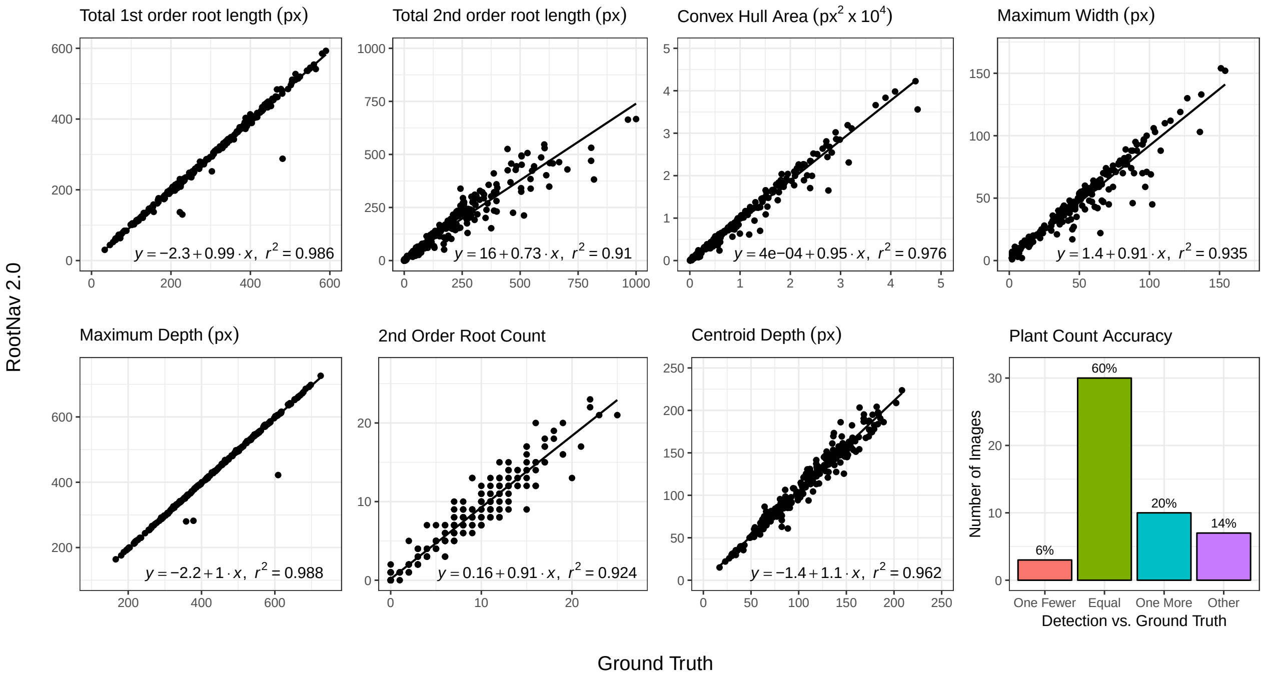


**Figure S1.** Extended plots showing quantitative results on the Arabidopsis Thaliana dataset of Rootnav 2.0 against the ground truth predictions for a variety of traits.

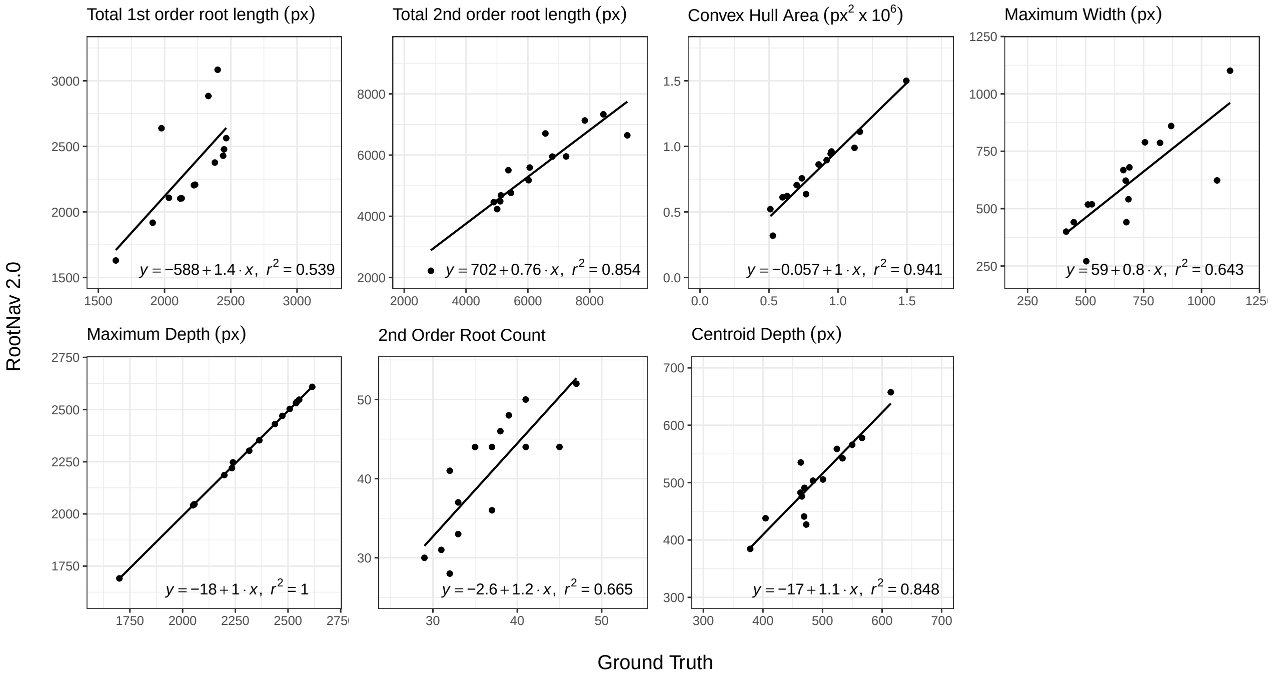


**Figure S2.** Extended plots showing quantitative results on the Brassica Napus dataset of Rootnav 2.0 against the ground truth predictions for a variety of traits.

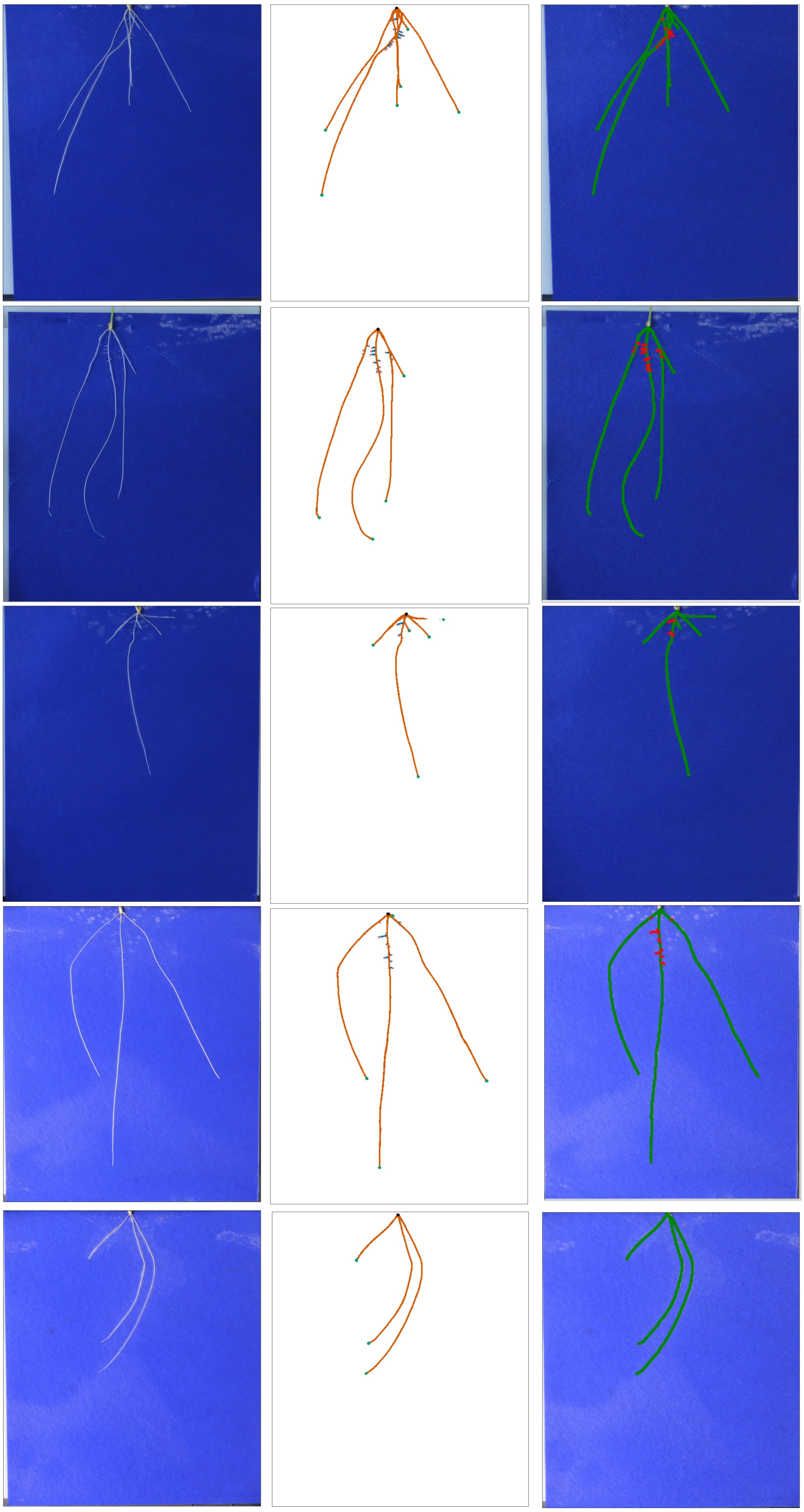

**Figure S3.** Example output of RootNav 2.0 on the wheat dataset. (Left) Original image. (Centre) Segmentation output showing root classes, tips and seeds. (Right) Original image overlaid with the reconstructed root system.

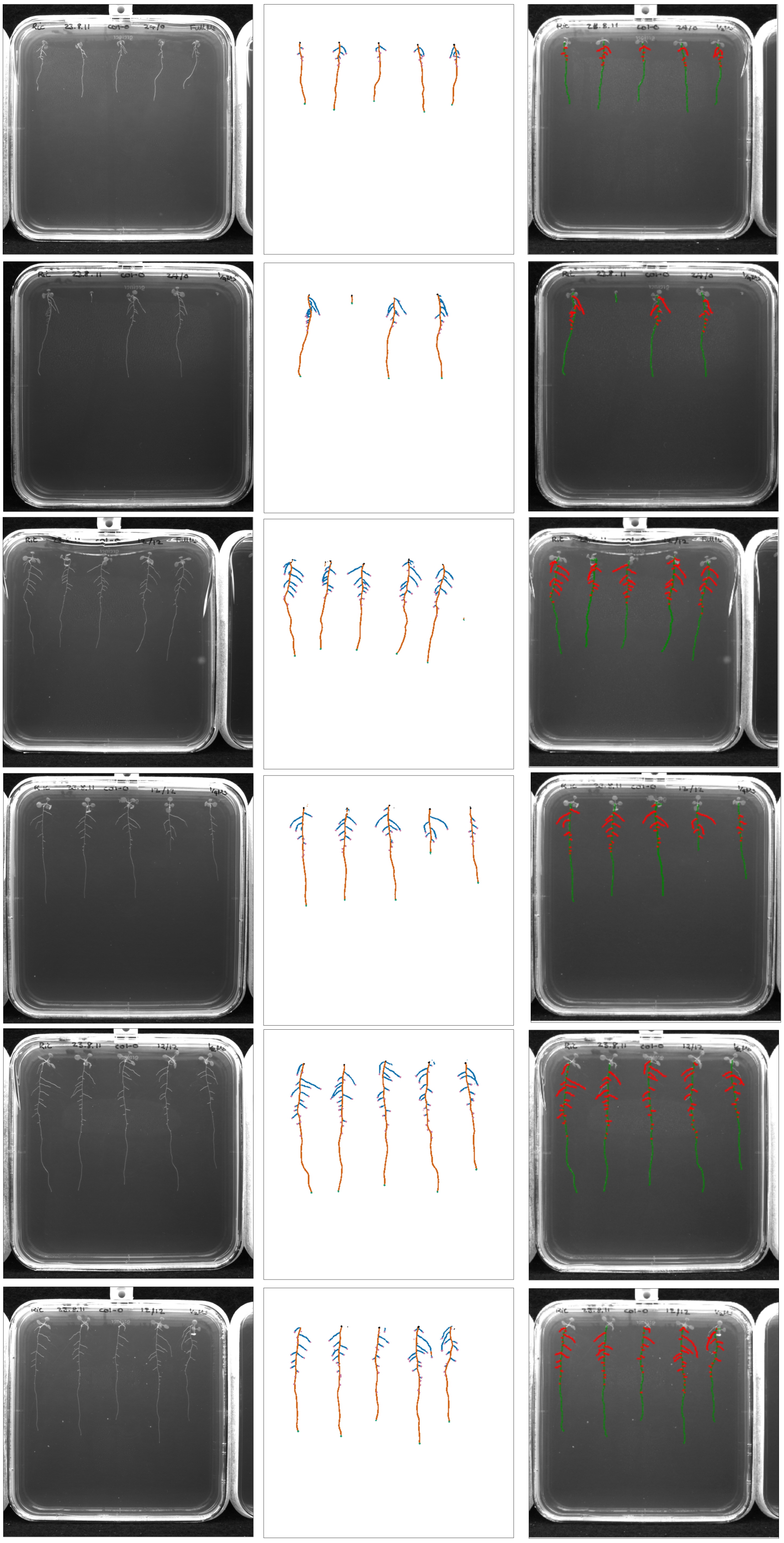

**Figure S4.** Example output of RootNav 2.0 on the Arabidopsis Thaliana dataser. (Left) Original image. (Centre) Segmentation output showing root classes, tips and seeds. (Right) Original image overlaid with the reconstructed root system.

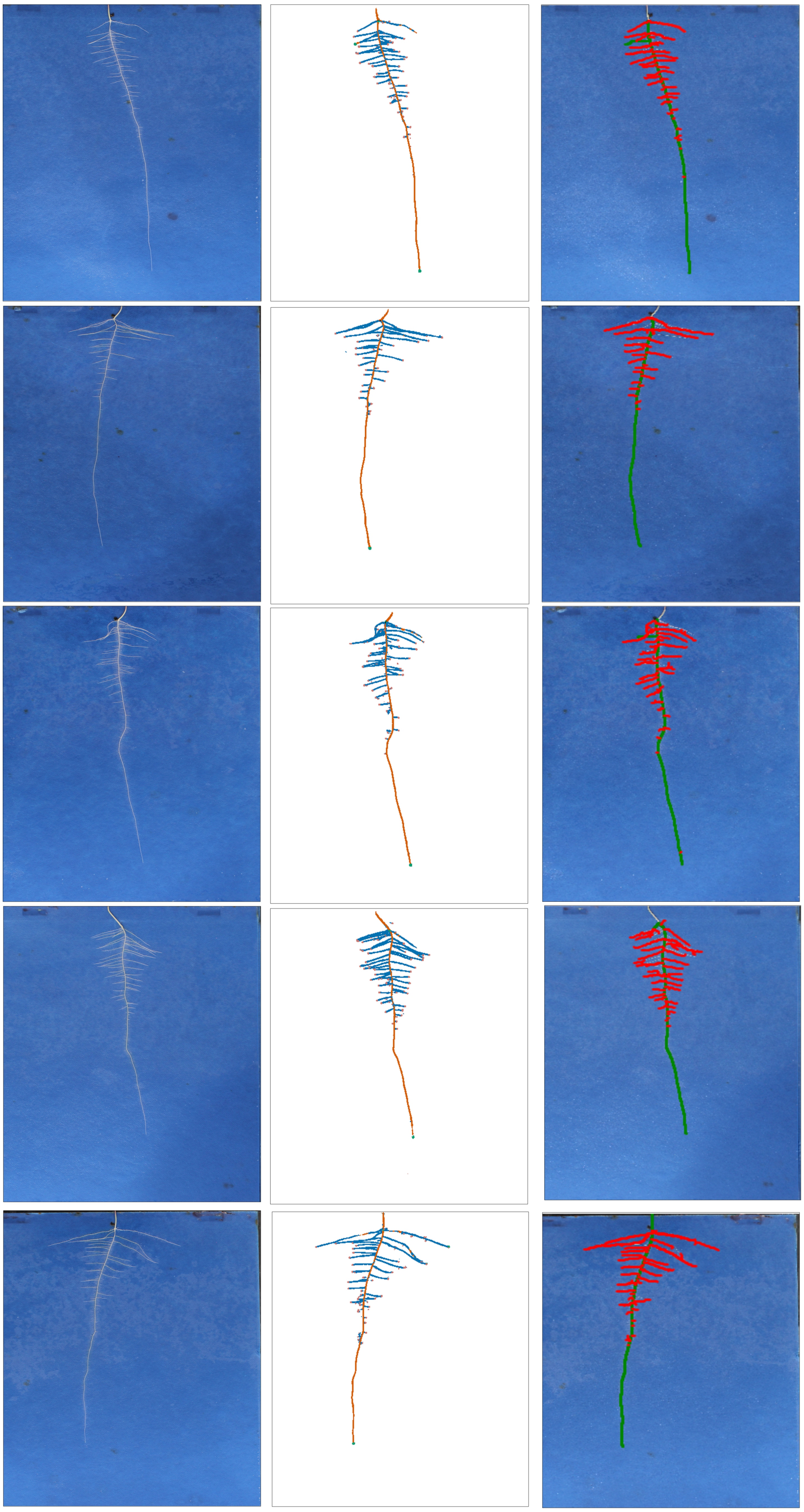

**Figure S5.** Example output of RootNav 2.0 on the Brassica Napus dataset. (Left) Original image. (Centre) Segmentation output showing root classes, tips and seeds. (Right) Original image overlaid with the reconstructed root system.

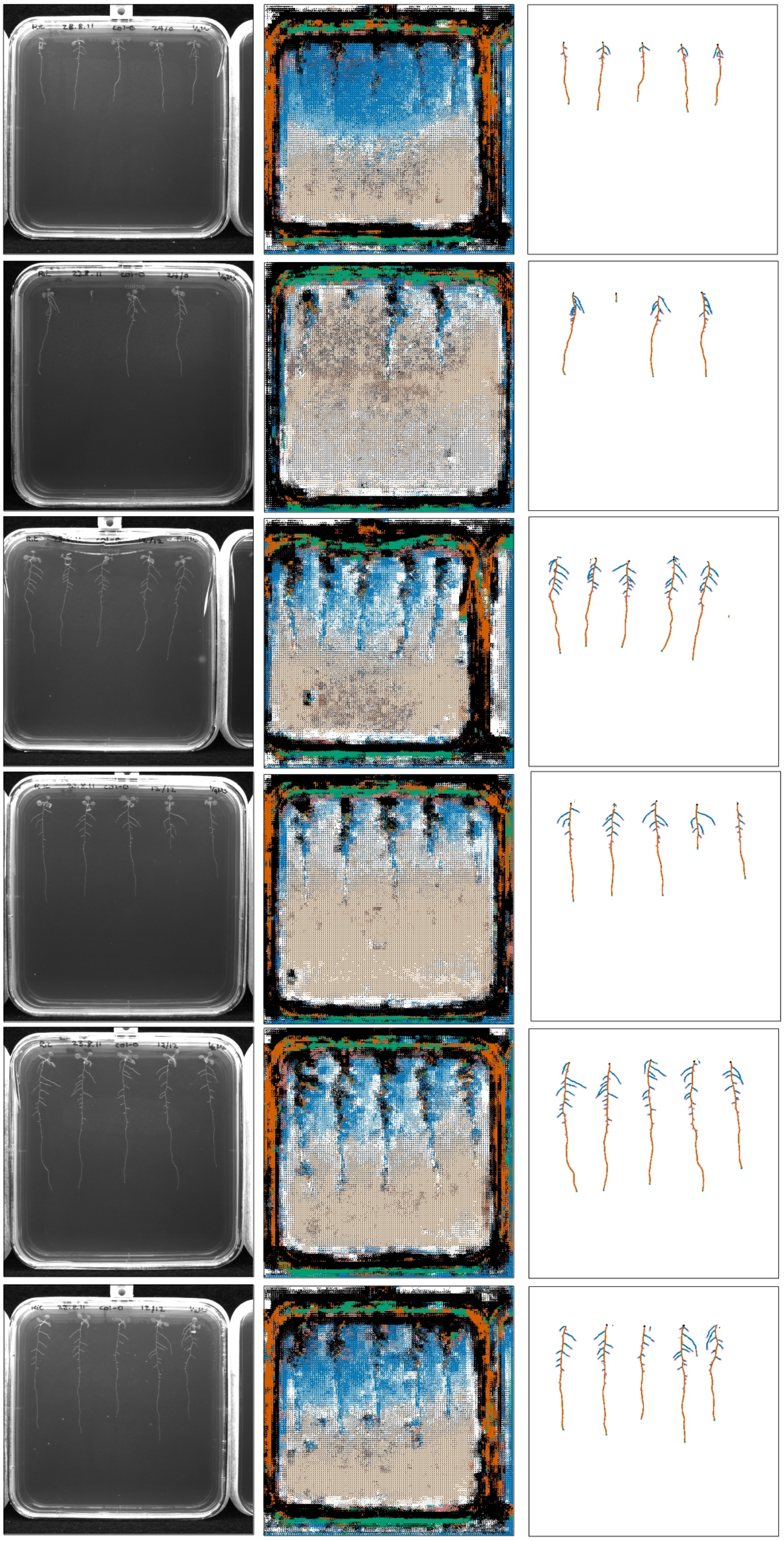

**Figure S6.** Example output on the *Arabidopsis* dataset showing the results of transfer learning vs. the results when the network is trained from scratch (randomly initialised weights). On this small dataset the network trained from scratch is unable to learn an adequate model. (Left) Original image. (Centre) Segmentation output for the network trained from scratch. (Right) Segmentation output of the trained network first initialised on the wheat dataset.

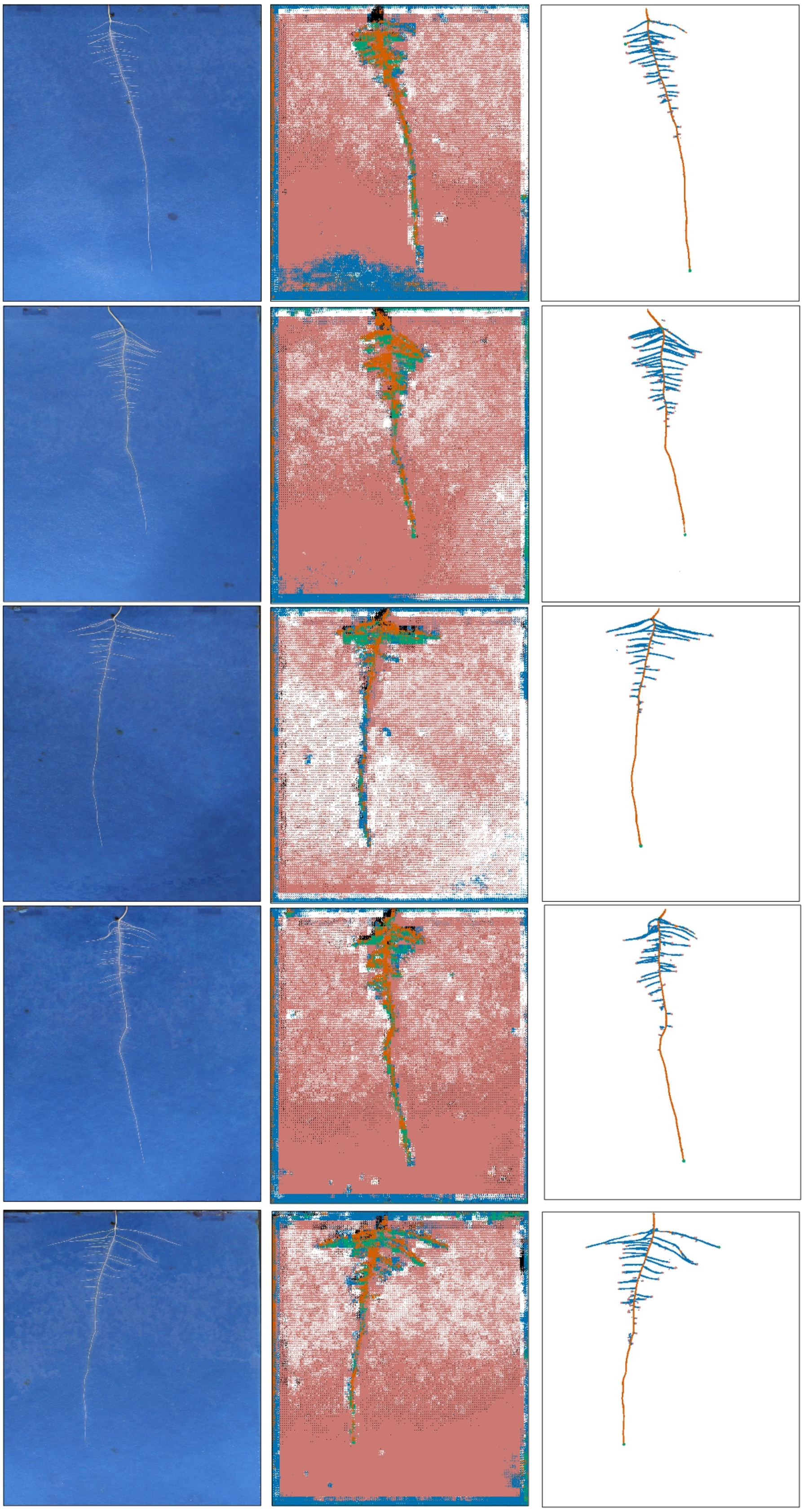

**Figure S7.** Example output on the rapeseed dataset showing the results of transfer learning vs. the results when the network is trained from scratch (randomly initialised weights). On this small dataset the network trained from scratch is unable to learn an adequate model. (Left) Original image. (Centre) Segmentation output for the network trained from scratch. (Right) Segmentation output of the trained network first initialised on the wheat dataset.
